# Dynamics of SARS-CoV-2 host cell interactions inferred from transcriptome analyses

**DOI:** 10.1101/2021.07.04.450986

**Authors:** Lukas Adam, Megan Stanifer, Fabian Springer, Jan Mathony, Chiara Di Ponzio, Roland Eils, Steeve Boulant, Dominik Niopek, Stefan M. Kallenberger

**Author notes:** these authors contributed equally to this work (Lukas Adam, Megan Stanifer).

## Abstract

The worldwide spread of severe acute respiratory syndrome-related coronavirus-2 (SARS-CoV-2) caused an urgent need for an in-depth understanding of interactions between the virus and its host. Here, we dissected the dynamics of virus replication and the host cell transcriptional response to SARS-CoV-2 infection at a systems level by combining time-resolved RNA sequencing with mathematical modeling. We observed an immediate transcriptional activation of inflammatory pathways linked to the anti-viral response followed by increased expression of genes involved in ribosome and mitochondria function, thus hinting at rapid alterations in protein production and cellular energy supply. At later stages, metabolic processes, in particular those depending on cytochrome P450 enzymes, were downregulated. To gain a deeper understanding of the underlying transcriptional dynamics, we developed an ODE model of SARS-CoV-2 infection and replication. Iterative model reduction and refinement revealed that a negative feedback from virus proteins on the expression of anti-viral response genes was essential to explain our experimental dataset. Our study provides insights into SARS-CoV-2 virus-host interaction dynamics and facilitates the identification of druggable host pathways supporting virus replication.

## Introduction

The novel coronavirus SARS-CoV-2 is responsible for the worldwide coronavirus disease 2019 (COVID-19) pandemic. Besides the respiratory epithelium of the nasopharynx and the lung, SARS-CoV-2 can infect different tissues of the human body including the mucosa of the intestine, the renal epithelium as well as lymphoid tissues (Gupta *et al*, 2020; Stanifer *et al*, 2020). Recent studies based on transcriptomics and proteomics techniques revealed host cell pathways affected by SARS-CoV-2, characterized molecular interactions between virus-host protein interactions, identified host factors potentially serving as therapeutic targets and developed strategies for drug repurposing (Gordon *et al*, 2020a, 2020b; Hadjadj *et al*, 2020; Bojkova *et al*, 2020; Klann *et al*, 2020; Blanco-Melo *et al*, 2020; Bouhaddou *et al*, 2020; Samavarchi-Tehrani *et al*, 2020; Schmidt *et al*, 2021; Selkrig *et al*, 2021; Stukalov *et al*, 2021; Triana *et al*, 2021; Wyler *et al*, 2021). Analyses of the translatome and proteome in human colorectal adenocarcinoma cells (Caco-2) showed that SARS-CoV-2 strongly affects translation, splicing, carbon metabolism and nucleic acid metabolism (Bojkova *et al*, 2020; Klann *et al*, 2020). A multi-omics study in a lung-derived cell line investigated virus-host protein interactomes and revealed that SARS-CoV-2 caused dysregulation of the TGF-β and EGFR pathways as well as autophagy (Stukalov *et al*, 2021). Investigations of molecular interactions between virus and host proteins suggested immediate interactions with cellular pathways involved in inflammation such as NF-κB, interferon or mTOR signaling (Gordon *et al*, 2020b; Triana *et al*, 2021). It was observed, for instance, that virus protein interactions with IL17RA immediately influence IL17 signaling. Virus proteins also directly interact with the mitochondrial outer membrane protein Tom70 that is involved in mitochondrial anti-viral signaling, through interaction with Hsp90, thereby affecting interferon signaling and apoptosis induction (Gordon *et al*, 2020a; Jiang *et al*, 2020; Stukalov *et al*, 2021; Samavarchi-Tehrani *et al*, 2020). SARS-CoV-2 inhibits the type I interferon response to evade the cellular anti-viral defense (Blanco-Melo *et al*, 2020; Chu *et al*, 2020; Triana *et al*, 2021; Stukalov *et al*, 2021). Analyses of phospho-proteomes further revealed that kinases involved in growth factor receptor signaling become activated upon SARS-CoV-2 infection (Bouhaddou *et al*, 2020; Klann *et al*, 2020; Stukalov *et al*, 2021). While kinases of the p38 pathway associated with inflammatory signaling were also triggered by SARS-CoV-2, Rho-associated protein kinases affecting organization of the interface between cytoskeleton and the plasma membrane were, in turn, inhibited. Moreover, phosphorylation of RNA-processing proteins was observed, which could indicate a strategy of the virus to prioritize translation of virus proteins over host cell proteins (Bouhaddou *et al*, 2020). Apart from affecting host proteins via specific interactions, SARS-CoV and SARS-CoV-2 both perturb the integrity of host cells through fragmentation of the Golgi apparatus and formation of viral replication organelles consisting of double-membrane vesicles tethered to the endoplasmic reticulum (Knoops *et al*, 2008; Snijder *et al*, 2020; Cortese *et al*, 2020).

Previous studies revealed several aspects of the host cell response on a qualitative level. However, a fine-grained temporal and quantitative analysis of the host cell response, in particular, in the early phase of SARS-CoV-2 infection, would be important and necessary for understanding SARS-CoV-2 infection at a systems level. Here, we developed a systems biology approach for studying transcriptional dynamics based on time-resolved RNA sequencing (RNA-seq) experiments analyzed using a set of expression profile functions fitted to all expressed genes. This enabled us to dissect the timing and interdepence of cellular processes that oppose or support virus replication. Using highly infectable Caco-2 (human colon cancer) cells as a model system, we observed a characteristic time-pattern of transcriptional upregulation associated with pathways involved in inflammation, kinase signaling, and processes related to cellular energy production, followed by transcriptional downregulation of various metabolic processes. We then developed a mathematical model of SARS-CoV-2 replication calibrated with experimental measurements and applied model reduction strategies to identify pathophysiologically required interactions between the virus and its host. Finally, we explored different strategies for direct and indirect interference with virus replication. Our study provides novel insights into the temporal pattern of host-virus interactions and can serve as a resource to develop strategies for combined targeting of cellular processes that support virus replication.

## Results

### SARS-CoV-2 rapidly reprograms its host

The dynamics of SARS-CoV-2 infection at the cellular level were studied using the model system of Caco-2 (human colorectal adenocarcinoma) cells. First, we compared two independent Caco-2 cell lineages. We observed striking differences in their infectivity which corresponded to the detected differences in ACE2 and cathepsin B expression (see Supplementary note S1, Supplementary Fig. S1). Using the high ACE2/cathepsin B Caco-2 lineage, we then infected cells at a multiplicity of infection (MOI) of 5 and collected RNA samples at eight time points between 0 and 48 hours post infection (hpi) followed by RNA-seq (Fig. 1A). In parallel, we detected cells expressing SARS-CoV-2 N protein by immunostaining (Fig. 1B). We observed that virus transcripts rapidly increased and peaked already at 12 hpi, amounting to about one third of all detected transcripts (Fig. 1C). Thereafter, virus transcripts decreased to about 17% of detected transcripts at 48 hpi. Read counts normalized to lengths of coding sequences (CDS) reflected the nested RNA architecture of nidovirales (Fig. 1D and Supplementary Fig. S2). Virus N proteins were detected in about one third of cells by immunostaining already at 4 hpi and the number of N protein positive cells increased to 100% within only 24 hours (Fig. 1E). Concurrently, the N protein expression levels per cells also increased (Fig. 1B). Cell counts remained constant until 24 hpi and decayed thereafter due to host cell death (Fig. 1F). The timing of virus release was aligned to the timing of host cell death, suggesting that virus particles were mostly released through cell lysis rather than budding from the plasma membrane of living cells (Fig. 1G). Taken together, SARS-CoV-2 replication was remarkably fast, resulted in high virus transcript loads already at 12 hpi and cell death, hence reflecting the typical characteristics of lytic viruses.

**Figure 1.**
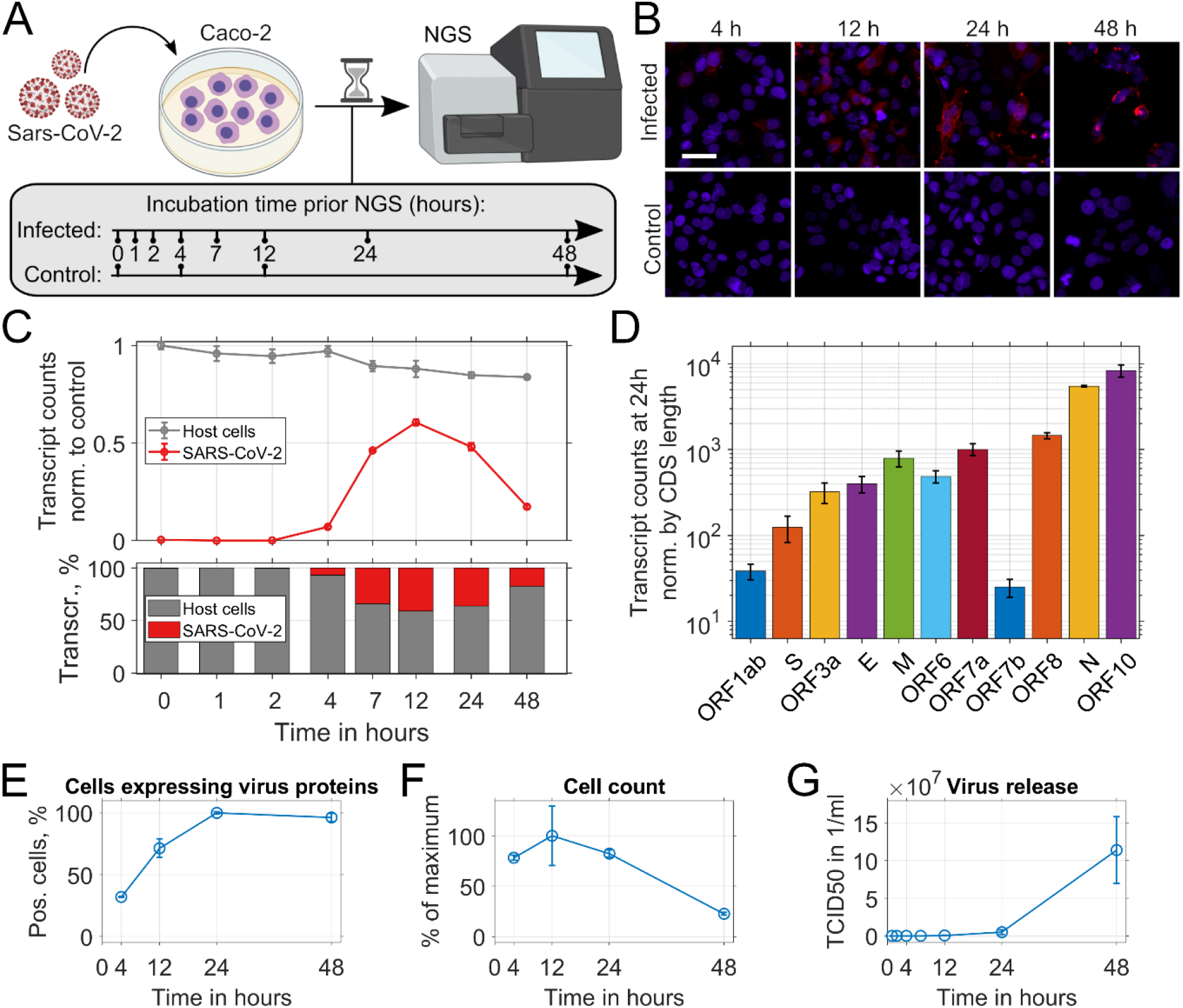
Dynamics of SARS-CoV-2 genome expression and virus replication. (**A**) Experimental setup. Caco-2 cells were infected with SARS-CoV-2 at an MOI of 5. Cells were lysed and RNA-seq was performed at the indicated time-points post infection. (**B**) Microscopy analysis of SARS-CoV-2 replication in Caco-2 cells infected with SARS-CoV-2 (top) or non-infected control cells (bottom). Representative images of three biological replicates are shown (blue, DAPI staining of nuclei; red, immunostaining of SARS-CoV-2 N protein; scale bar, 50 μm). (**C**) Analysis of absolute (top) and relative (bottom) host cell and virus transcript counts. At 12 hpi, virus transcripts peaked, constituting 41% of all transcripts (means of n=3 replicates; error bars, SEM). (**D**) Virus transcript read counts increased from ORF1 to ORF10, reflecting the nested RNA architecture of SARS-CoV-2 (bars: average read counts of n=3 replicates normalized by CDS length at 24 hpi; error bars, SEM; CDS, coding sequence). (**E**, **F**) Microscopy analysis of the fraction of cells with detectable expression of SARS-CoV-2 N protein (**E**) and the normalized total cell count (**F**) at the indicated time points.(**G**) Quantification of the released virus particles by endpoint dilution assay (TCID50, 50% tissue culture infective dose). (E-G) n=3, error bars indicate standard deviation.

### SARS-CoV-2 infection triggers a distinct sequence of transcriptional responses in host cells

To characterize the transcriptional response in host cells upon SARS-CoV-2 infection, we analyzed changes in transcriptomes over time, first at the macroscopic level. Distributions of log2 fold changes of gene expression (L) indicated a fraction of strongly upregulated genes (defined by L ≥ 1) that was highest between 4 and 12 hpi (Fig. 2A, at maximum 6.54%). In contrast, the fraction of genes for which expression was strongly downregulated (defined by L ≤ − 1) steadily increased between 4 hpi and 48 hpi (Fig 2A, at maximum 4.15%). We conclude that the host response shows a characteristic, rapid upregulation in a fraction of genes within the first few hpi, followed by a phase of downregulation towards later time points.

**Figure 2.**
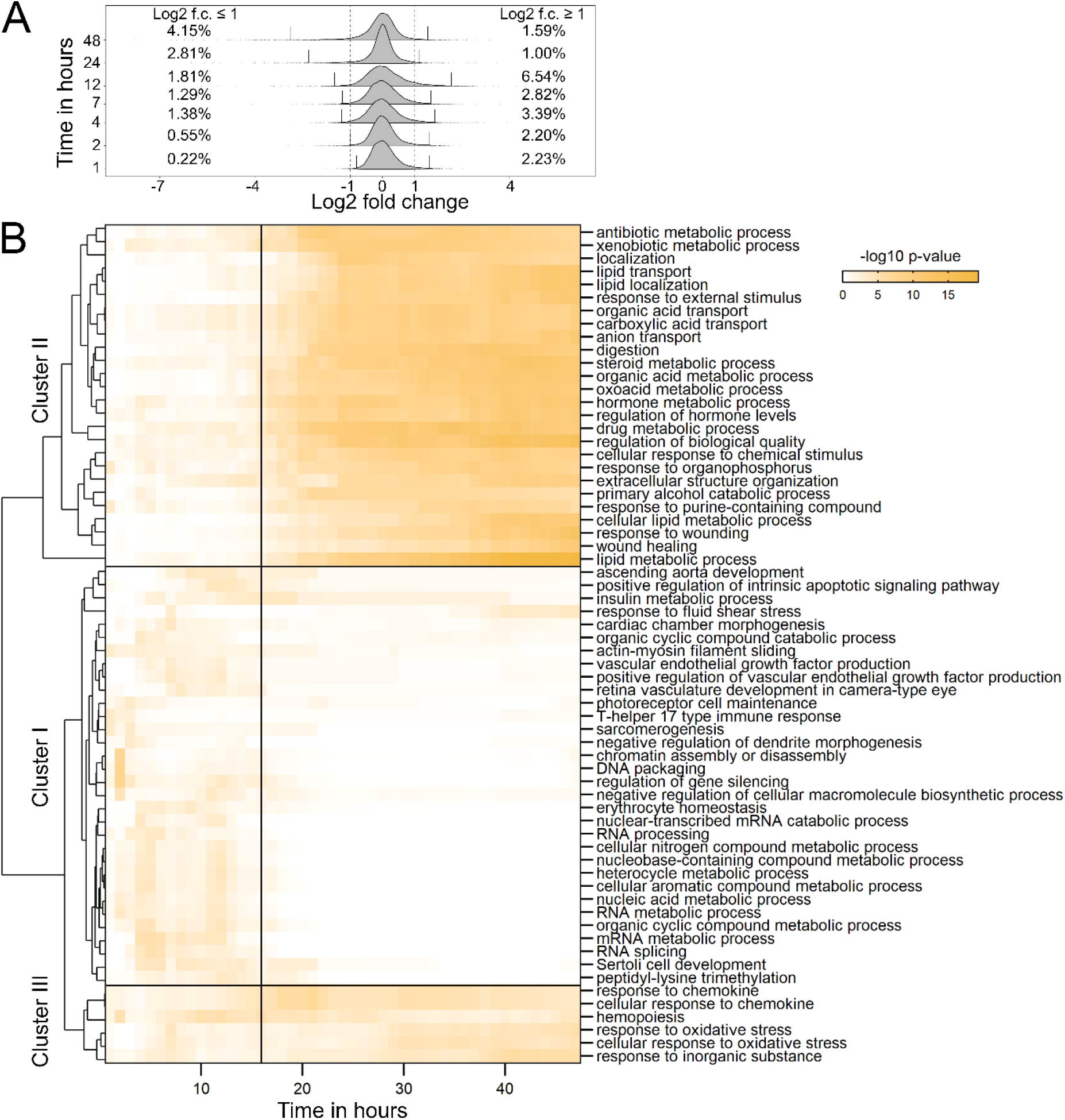
Dynamic alteration of the host cell transcriptome upon SARS-CoV-2 infection. (**A**) Distributions of log2 fold changes of normalized and background-corrected gene transcript reads relative to initial transcript counts, and percentages of genes with log2 fold changes *L* ≥ 1 or *L* ≤ −1 (dashed lines, log2 fold changes of ±1; vertical lines, 2.5 and 97.5 percentiles). (**B**) Clusters of p-values from GO term enrichment at interpolated time intervals indicate the dynamics of cellular processes in response to SARS-CoV-2 infection. The following qualitative patterns can be distinguished: early, transient response defined by transient expression changes within 12 hpi (cluster I), late, sustained response characterized by expression changes that take place after 12 hpi and are maintained (cluster II), and an early, sustained response indicated by maintained expression changes that already start at few hours post infection (cluster III).

Next, to identify cellular processes underlying these dynamics, the time series of log2 fold changes in transcript counts were subjected to gene ontology (GO) term enrichment analysis in interpolated hourly time intervals. We then clustered time courses of p-values indicating GO term enrichment. Three qualitatively different patterns of early, late or continuously regulated cellular processes were observed (Fig. 2B). Cluster I, the cluster overrepresented within the first 12 hpi, contained several GO terms associated with mRNA biogenesis such as ‘nuclear-transcribed mRNA catabolic process’, ‘RNA processing’, ‘RNA metabolic process’ or ‘RNA splicing’ (Fig. 2B, cluster I). Cluster II, the cluster overrepresented in late time points of infection, contained various GO terms involved in cellular metabolism including the terms ‘steroid metabolic process’, ‘organic acid metabolic process’, ‘hormone metabolic process’ as well as processes involved in the cellular response to drugs and chemical agents such as ‘antibiotic metabolic process’, ‘xenobiotic metabolic process’, ‘drug metabolic process’ or ‘cellular response to chemical stimulus’. Cluster III, the cluster which represents continuously regulated processes, contained GO terms involved in cytokine signaling and the cellular stress response. These results suggest that upon SARS-CoV-2 infection, a rapid transcriptional response takes place well before high virus transcript levels are detectable (Fig. 1C). The observed transcriptional regulation can be separated into cellular processes already influenced in the first hours after infection, delayed effects appearing after 12 hpi, i.e. when the amount of virus transcripts already starts to decrease, and gradually affected cellular processes.

### Dissecting the dynamics of virus-host interactions using an expression profile function approach

Next, we analyzed temporal sequences and amplitudes of transcriptional changes associated with cellular processes and signal transduction pathways. Taking into account that cellular processes were continuously or transiently affected, we fitted four profile functions to log2 fold changes in all *N* = 13,322 expressed genes. The functions described a continuous or transient increase or decrease in gene expression (Fig. 3A). Profile function fits were then used to extract time points of expression changes, defined by the times at half-maximal increase or decrease, as well as amplitudes. To focus on curated graphs of signaling pathways, we analyzed transcriptional effects mapped to the KEGG database (Kanehisa & Goto, 2000; Kanehisa *et al*, 2019). To visualize temporally ordered expression changes in cellular processes, we then created time-resolved pathway charts of strongly affected genes defined by absolute values of log2 fold changes |*L*| ≥ 1. Fig. 3B–E visualizes the time points of maximal expression changes for genes of the pathway ‘Transcription factors’, as extracted from profile function fits (Fig. 3A). In this visualization, borders of colored stripes indicate turning points of expression profile functions (see Supplementary Fig. S3 for further pathway charts).

**Figure 3.**
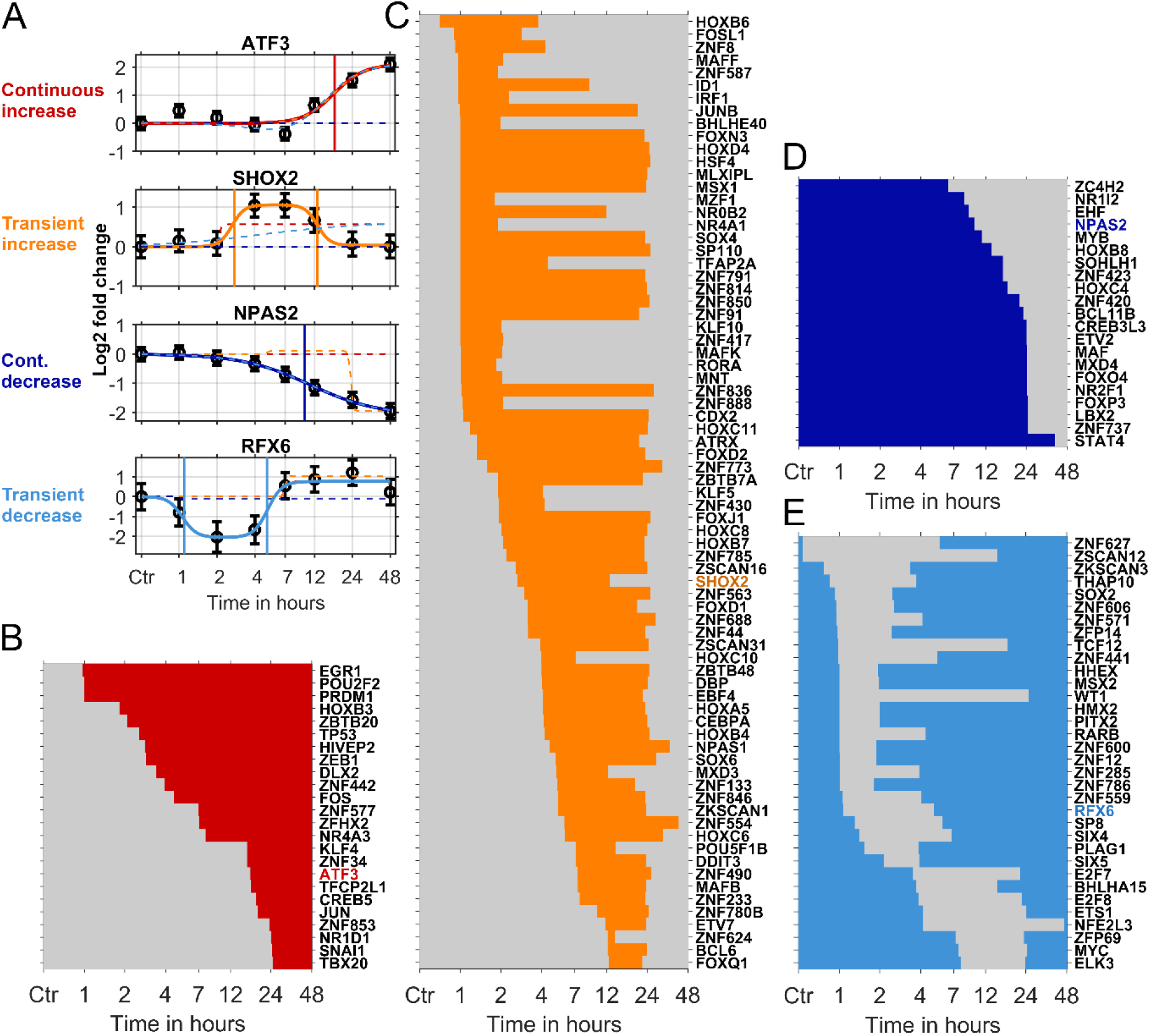
Transcriptionally altered host genes show distinct dynamic patterns. (**A**) According to the observed transcription dynamics, four functions describing either continuous or transient increase (red, yellow) or decrease (dark or light blue) were fitted to transcription fold changes of all detected genes. Optimal function types were selected based on smallest BIC values (solid lines, function types with smallest BIC values; dashed lines, fits of other function types). Points of expression changes, indicated by vertical lines, represent the time points at half maximal increase or decrease according to the extracted functions. (**B**–**E**) To analyze the transcriptional dynamics of the host cell response, time points of expression changes (up- or downregulation) of genes associated with KEGG pathways as ‘Transcription factors’ were displayed as lines indicating the time interval of increased transcription. (**B**) Continuous increase, (**C**) transient increase, (**D**) continuous decrease, (**E**) transient decrease. Genes shown in **A** were highlighted in respective profile function colors.

To identify strongly affected cellular processes and study the temporal order of transcriptional responses connected to these processes, we determined absolute numbers and fractions of strongly up- or downregulated genes, defined by |*L*| ≥ 1, in all KEGG pathways (Fig. 4). Pathways with largest absolute numbers were ‘Transcription factors’, ‘Membrane trafficking’, ‘Chromosome and associated proteins’, ‘Exosome’ as well as ‘Peptidases and inhibitors’ (Fig. 4A). Consistent with our analysis based on GO terms, the pathways ‘Cytochrome P450’ and ‘Cytokines and growth factors’ were among those with largest fractions of strongly affected genes (Fig. 4B). To analyze the timing of effects in pathways, we extracted time points, at which 50% of the affected genes were regulated. To this end, cumulative sums of time intervals with increased expression of strongly regulated genes were determined. The top 25 up- and down-regulated pathways, with the largest fractions of strongly regulated genes were then sorted according to times when 50% of genes were affected (Fig. 4C, D). We observed an early response in several pathways associated with inflammation and cytokine signaling (e.g., ‘Viral protein interaction with cytokine and cytokine receptors’, ‘IL-17 signaling pathway’, ‘TNF signaling pathway’, ‘Cytokine-cytokine receptor interaction’) already around 1 hpi (Fig. 4C). Sets of genes upregulated in ‘IL-17 signaling pathway’, ‘TNF signaling pathway’ or ‘Cytokine-cytokine receptor interaction’ showed large intersections with other pathways involving kinase signaling as ‘MAPK signaling pathway’ or ‘PI3K-Akt signaling pathway’ (Supplementary Fig. S3A–E). Thereafter, around 2 hpi, several processes related to translation and production of chemical energy in mitochondria became upregulated, such as ‘Ribosome’, ‘Thermogenesis’ or ‘Oxidative phosphorylation’ (Fig. 4C). Several pathways ranked among those with largest fractions of strongly upregulated genes because they contain many genes involved in mitochondria or ribosomes (green and blue font in Fig. 4C; pathway charts for ‘Ribosome’ and ‘Thermogenesis’ in Supplementary Fig. S3G, H). Among these was the pathway ‘Coronavirus disease – COVID-19’ (Supplementary Fig. S3Q) that contains various genes involved in ribosome function. Interestingly, all of these pathways were upregulated before virus transcript levels strongly increased (Fig. 4C, top panel). In most cases, upregulation was reverted between 24 and 48 hpi, i.e. when cells died and virus particles were released (Fig. 1F, G).

**Figure 4.**
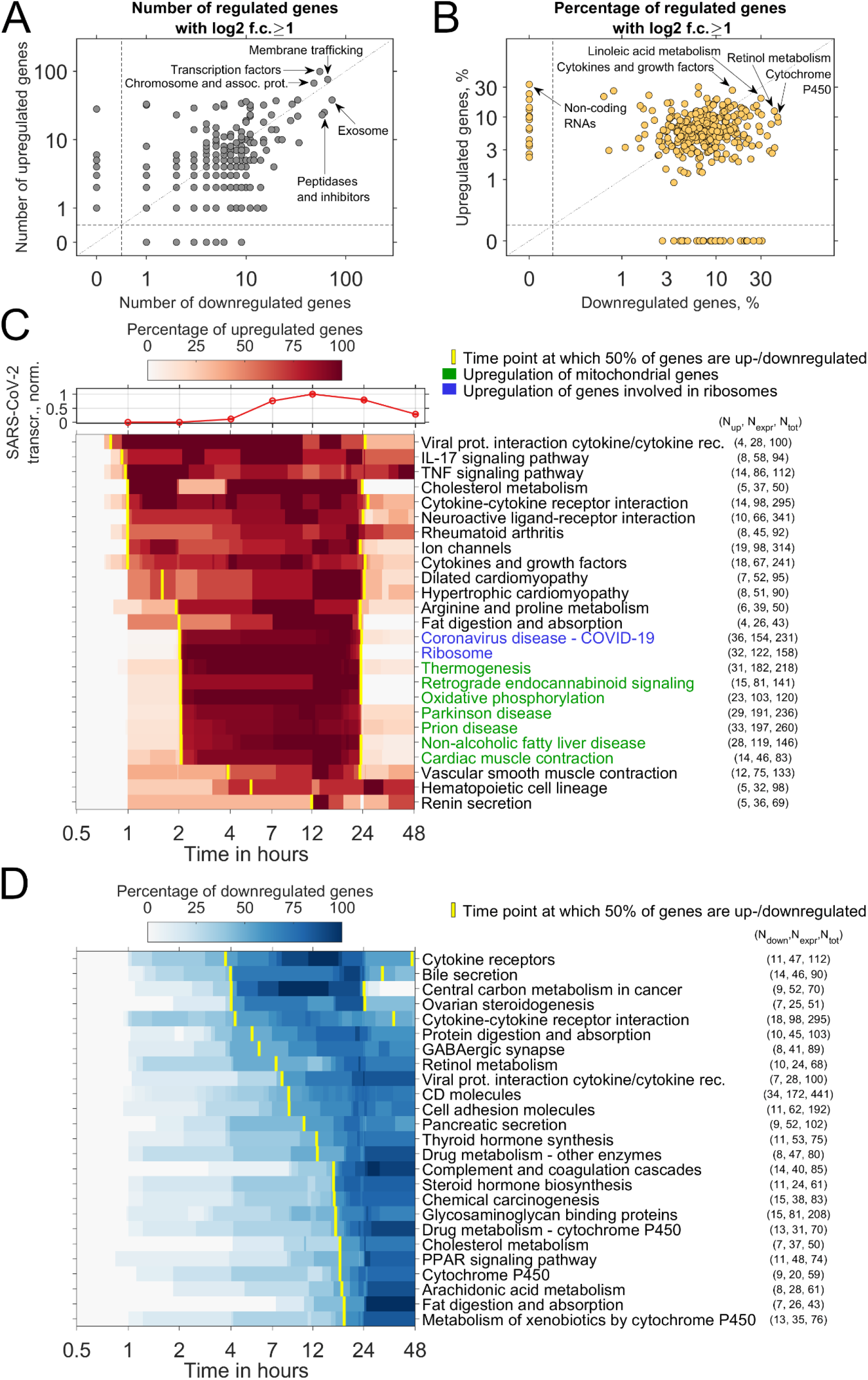
Transcriptional dynamics is associated with selected cellular pathways. (**A**) Scatter plot showing the number of regulated genes in KEGG pathways affected by SARS-CoV-2 infection. Pathways comprising strongly up- or downregulated genes, defined by log2 fold changes of at least 1, were selected. (**B**) Scatter plot of KEGG pathways affected by SARS-CoV-2 infection. Dots indicate the percentage of strongly regulated genes relative to the total number of expressed genes within a pathway. (**C**) 25 cellular pathways with the largest fractions of upregulated genes, selected from pathways with at least 10 strongly regulated genes. Pathways were sorted according to times at which 50% of all strongly regulated genes were up- or downregulated (top to bottom). (**D**) Top 25 cellular processes with largest fractions of downregulated processes as in **C**. Transcription was downregulated, delayed relative to the observed upregulation, in various processes involved in metabolism (N_up/down_, number of up- or downregulated genes; N_exp_, number of expressed genes detected by RNA sequencing; N_tot_, total number of genes within KEGG pathways).

The top 25 pathways with largest fractions of strongly downregulated genes showed that the gene expression mostly declined at later time points (Fig. 4D). Specifically, two phases of downregulation could be observed, a first phase between 4 and 12 hpi, i.e. when virus transcript levels peaked, and a second phase between 12 and 48 hpi, i.e. when virus transcript levels decreased (Fig. 4D). Around 4 hpi, the pathway ‘Cytokine receptors’ was downregulated subsequent to the early upregulation of several pathways depending on cytokines (compare Fig 4C and D). The initially upregulated pathways ‘Cytokine-cytokine receptor interaction’, ‘Viral protein interaction with cytokine and cytokine receptors’ and ‘Fat digestion and absorption’, were downregulated at times when large virus transcript levels were present between 4 and 24 hpi (Fig 4C and D). Strongly downregulated pathways included several cellular processes involved in metabolism such as ‘Protein digestion and absorption’, ‘Steroid hormone biogenesis’, ‘Cholesterol metabolism’ or ‘PPAR signaling pathway’. Strikingly, multiple pathways related to xenobiotic metabolism depending on cytochrome P450 enzymes were affected as part of the second phase of downregulation between 24 and 48 hpi (Fig. 4D). The observation that several unexpected pathways as ‘Bile secretion’ emerged from our analysis can be explained by intersections of genes involved in common metabolic processes. Large intersections of genes, for instance, were present between the pathways ‘Bile secretion’ and ‘Ovarian steroidogenesis’ with pathways related to xenobiotic metabolism, as well as between ‘Central carbon metabolism in cancer’ and ‘Cytokine receptors’. Several enzymes involved in the metabolism of specific drugs, including CYP2B6, CYP2C19 or CYP3A5 were downregulated (‘Drug Metabolism – cytochrome P450’ pathway, Supplementary Fig. 3K). The pathway ‘Complement and coagulation cascades’, that was downregulated mostly between 12 and 48 hpi, contains several protease inhibitors from the serpin superfamily, such as SERPINA1 (α1-antitrypsin), SERPINA5 (protein C inhibitor), SERPINC1 (antithrombin III), SERPIND1 (heparin cofactor II) or SERPINF2 (α2-plasmin; Supplementary Fig. S3J). Of these, SERPINA1 and SERPINA5 serve as inhibitors of the host-factor proteases for SARS-CoV-2, TMPRSS2 and cathepsin L (Fortenberry *et al*, 2006; Wettstein *et al*, 2021). Further serpins, such as SERPINB8, an inhibitor of the host-factor protease furin (Dahlen *et al*, 1998), were downregulated as part of the pathway ‘Peptidases and inhibitors’ (Supplementary Fig. 3R). Moreover, we observed downregulation of ACE2 as part of the ‘Peptidases and inhibitors’ pathway, mirroring previous findings in SARS-CoV-infected Vero cells and human intestinal organoids (Glowacka *et al*, 2010; Stanifer *et al*, 2020).

In summary, we observed a sequence of early upregulation of transcription in pathways related to inflammation and cytokine signaling followed by upregulation in pathways associated with translation and chemical energy production between 1 and 4 hpi. Eventually, several pathways involved in cellular metabolism, particularly in xenobiotic metabolism, were strongly downregulated.

### Dynamic model of SARS-CoV-2 replication and the anti-viral response

To obtain mechanistic insights to the dynamics of SARS-CoV-2 replication in conjunction with the anti-viral response in host cells, we developed mathematical models consisting of coupled ordinary differential equations. We aimed at finding a parsimonious model, with minimal complexity, to identify interactions required for explaining our experimental dataset. In the optimal model version, virus transcripts *V* evoke transcription of mRNAs of anti-viral response genes *m_A_* (Fig. 5A). These are translated to anti-viral proteins *A* that inhibit virus replication. This assumption of a direct interference with virus replication was motivated by the evidence that host-cell proteins that are part of the anti-viral response cause nonsense-mediated mRNA decay (NMD) in SARS-CoV-2 and other coronaviruses (Wada *et al*, 2018; Schmidt *et al*, 2021). Virus transcripts are then translated to virus proteins *P* that inhibit transcription of anti-viral response genes. The model was calibrated with SARS-CoV-2 transcript counts, SARS-CoV-2 N protein measurements, and a cumulative function describing the expression of genes associated with the anti-viral response (Fig. 5B). This cumulative function was derived from time points at which expression of anti-viral response genes increased or decreased (Supplementary Fig. S4, see Methods for details). Virus transcripts were scaled to the total amount of cellular transcripts.

**Figure 5.**
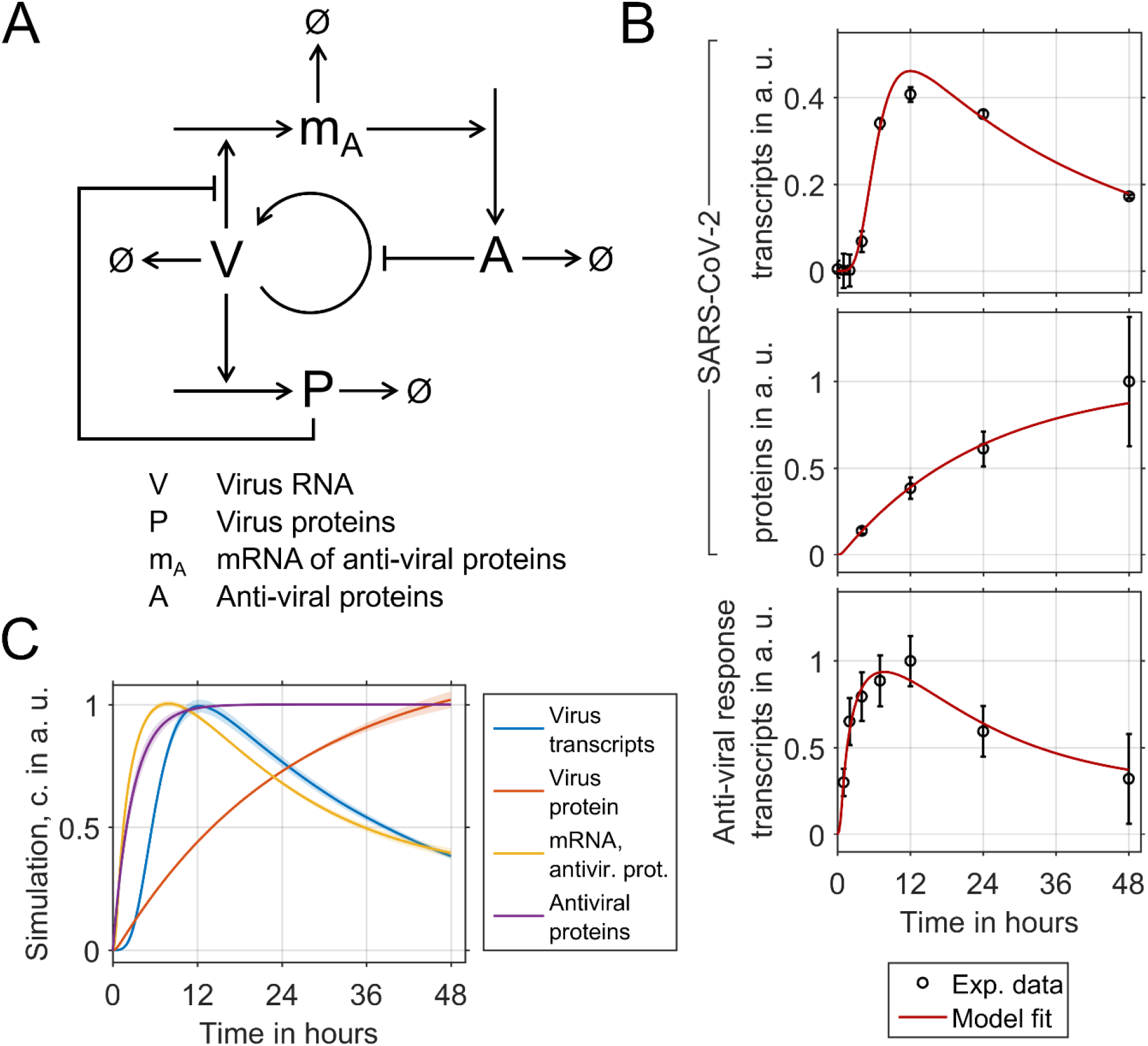
Dynamic model of SARS-CoV-2 replication and the anti-viral response. (**A**) Overview of the model describing virus replication, synthesis of virus proteins and expression of transcripts and proteins associated with the anti-viral response exhibited by the host cell. In the model, viral RNA (*V*) stimulate the expression of anti-viral response gene transcripts (*m_A_*) that are translated to proteins (*A*) that inhibit virus replication. Viral transcripts are translated to virus proteins (*P*) that inhibit translation of *m_A_*. (**B**) Best model fits to measurements of virus transcripts (top), virus N protein (center), and average expression levels of anti-viral response genes (bottom). (**C**) Model predictions of virus transcripts and proteins as well as mRNAs and proteins associated with the anti-viral response (lines, predictions using means of parameters from best 10 of 5000 fits; shaded areas, 1σ-confidence intervals; a. u., arbitrary units).

To identify processes required for explaining the experimental dataset and find an optimal model version, we iteratively extended an initial model and refitted model variants. Model extensions were pertained in case modified versions resulted in an improved model fit, assessed by values of the Bayesian information criterion (BIC; see Methods and Supplementary Figs. S5–7 for details, Supplementary Tables S1 and S2 for equations and parameter estimates). Several strategies used by SARS-CoV and SARS-CoV-2 to evade the host cell anti-viral response and perturb translation in host cells were previously described and translated to possible model extensions as described below. It was shown that SARS-CoV-2 inhibits the anti-viral interferon response at the transcriptional and proteome levels (Hadjadj *et al*, 2020; Stukalov *et al*, 2021). Previous studies also found that SARS-CoV actively inhibits host cell translation (Narayanan *et al*, 2008; Nakagawa *et al*, 2016) through the interaction of nsp1 and the 40S ribosomal subunit and inducing cleavage of host cell mRNAs (Kamitani *et al*, 2006, 2009; Huang *et al*, 2011). Furthermore, the nsp1 protein of SARS-CoV counteracts host cell translation by inhibiting the 48S initiation complex formation (Lokugamage *et al*, 2012).

Therefore, we tested whether including the following components improved the model, based on reductions in BIC: (1) negative feedback of *P* on the transcription of *m_A_*, (2) negative feedback of *P* by cleavage of *m_A_*, (3) negative feedback of *P* on the synthesis of *A*. Further, we tested, whether including (4) positive feedback of *P* on replication of *V*, or (5) positive feedback of *P* on virus replication by virus release and influx could improve the model. Based on iterative refinement and model selection from a total of 18 model variants, we found that it was indeed necessary to include the negative feedback of *P* on the transcription of *m_A_* to explain the experimental dataset (Supplementary Figs. S5–7). On the contrary, including the previously described cleavage of host-cell mRNAs by virus proteins, the inhibition of the translation of anti-viral proteins by virus proteins, or reactions describing positive feedback of *P* on virus replication did not result in further model improvement. Parameter estimates indicated that the production of virus proteins was already saturated at small total virus transcript levels, which suggests that the major part of virus transcripts was not required for producing actual virus particles. Further, the anti-viral response was already saturated at a small amount of virus transcripts. Interestingly, model simulations predicted that transcription of anti-viral genes as well as translation to anti-viral proteins was induced within 6 hpi before the peak level of virus transcripts at 12 hpi (Fig. 5C).

In conclusion, a mathematical model was developed that is capable of describing the dynamics of SARS-CoV-2 replication and the anti-viral response. Model discrimination suggested a strong inhibitory effect of SARS-CoV-2 proteins on the anti-viral response, and strong over-saturation of the translation of virus transcripts as well as the induction of anti-viral genes.

### Strategies for direct and indirect interference with SARS-CoV-2 replication

Subsequent to characterizing the cellular processes affected by SARS-CoV-2, we investigated two complementary strategies for interfering with virus replication. First, we developed an approach for experimentally characterizing direct inhibitors of SARS-CoV-2 protein maturation. Then, indirect inhibitors of host cell pathways presumably supporting virus replication were tested.

SARS-CoV-2 replication requires auto-catalytic cleavage of the precursor proteins pp1a and pp1ab by the 3C-like protease (3CL^pro^; also referred to as M^pro^), which represents a potential drug target for direct interference with virus replication. To monitor the activity of 3CL^pro^ and effects of possible protease inhibitors, we developed a live-cell assay based on fluorescent cleavage probes. These probes comprised a fusion of a nuclear export sequence (NES), a peptide sequence specifically cleaved by 3CL^pro^ and GFP (Fig. 6A). Before cleavage, probes are restricted to the cytoplasm. Following cleavage-mediated removal of the NES, GFP can enter the nucleus and equilibrates between nucleus and cytoplasm. Probe cleavage in single cells can hence be followed in real time by confocal live-cell imaging and quantified using the GFP fluorescence intensity in the nucleus *I_ncl_* and cytoplasm *I_cpl_*. We assumed that the concentration of intact probes in the cytoplasm [*PrF*] is proportional to the difference of fluorescence intensities [*PrF*]~(*I_cpl_* − *I_ncl_*).

**Figure 6.**
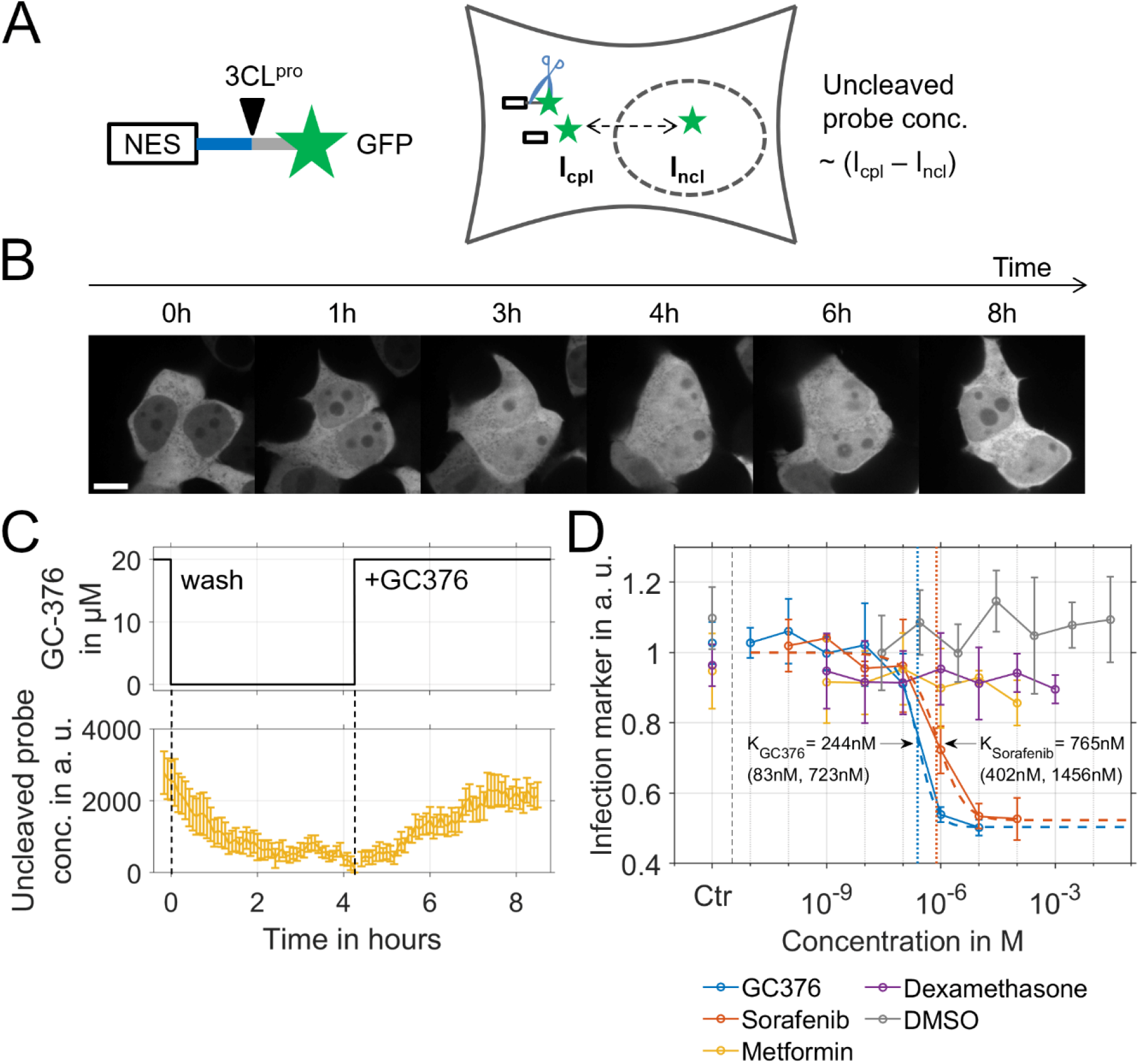
Direct and indirect inhibition of SARS CoV-2 replication. (**A**) Experimental setup. Cleavage probes consisting of a nuclear export sequence (NES), a 3CL^pro^ cleavage site (NS4-NS5) and GFP were expressed in HEK293T cells. Uncleaved probes are continuously exported from the nucleus and hence located in the cytoplasm. Following 3CL^pro^-mediated cleavage, GFP enters the nucleus by diffusion. The cytoplasmic and nuclear fluorescence intensities (I_cpl_, I_ncl_) were used to calculate the concentration of the uncleaved probe. (**B**, **C**) HEK293T cells co-expressing a cleavage probe containing the cleavage site between NSP4 and NSP5 and 3CL^pro^ were incubated in presence of 20 mM GC376, the drug was removed at time point 0 and re-added at 250 minutes (scale bar, 10 μm). Representative microscopy images (**B**) and corresponding quantification of the concentration of the uncleaved probe over time (**C**). Nuclear signal increase following GC376 removal indicates probe cleavage (data in **C**,means of n=10 cells, error bars: SEM). (**D**) Caco-2 cells were infected in the presence or absence of inhibitors. Virus replication was assessed at 24 hpi via immunofluorescence of SARS-CoV-2 N protein (means of n=3 replicates, error bars: SEM).

Peptide sequences of all 11 cleavage sites processed by 3CL^pro^ were integrated in cleavage probes (Supplementary Table S1), which were co-expressed with 3CL^pro^ in HEK293T cells. Using GFP-immunoblotting, we found that all probes were cleaved albeit with varying efficiency (Supplementary Fig. S8). In line with recent studies, we observed that the broad-spectrum coronavirus protease inhibitor GC376 could prevent probe cleavage (Fig. 6B) (Vuong *et al*, 2020; Fu *et al*, 2020), as indicated by a predominant cytoplasmic fluorescence. After washing cells with medium lacking the inhibitor, the nuclear fluorescence increased indicating probe cleavage (Fig. 6B, C). Re-adding GC367 resulted in inhibition of 3CL^pro^ as indicated by a decrease of the nuclear and an increase of the cytoplasmic fluorescence intensity due to re-synthesis of intact probes and degradation of cleavage products. Collectively, the developed live-cell assay can be used for testing potential inhibitors of 3CL^pro^ and studying the kinetics of protease inhibition in single cells. Subsequently, we tested whether GC367 could inhibit SARS-CoV-2 replication in our experimental system (see below).

In addition to blocking SARS-CoV-2 polyprotein cleavage, we tested complementary strategies for indirect interference with virus replication. As shown by our transcriptomic analyses, SARS-CoV-2 infection results in rapid transcriptional upregulation in pathways involved in kinase signaling as the IL17, TNF, MAPK or PI3K/Akt pathways as well as processes involved in translation and production of chemical energy that likely support SARS-CoV-2 replication (Fig. 4C). The PI3K/Akt/mTOR and MAPK pathways control translation based on phosphorylation of eukaryotic translation initiation factors (eIFs) and associated regulators (Roux & Topisirovic, 2012). It was previously shown that activation of mTOR signaling regulates energy metabolism and stimulates mitochondrial activity as well as biogenesis thereby increasing the capacity for ATP production (Cunningham *et al*, 2007; Düvel *et al*, 2010; Morita *et al*, 2013). Based on this hypothesis, it is reasonable to assume that inhibiting these pathways might counteract virus replication. Therefore, we tested the effect of metformin, an inhibitor of the NF-κB and the PI3K/Akt/mTOR pathways (Dowling *et al*, 2007; Hattori *et al*, 2006; Kalender *et al*, 2010; Moiseeva *et al*, 2013; Green *et al*, 2010), and an inhibitor of mRNA translation in cancer cells (Zakikhani *et al*, 2006), on virus replication. Moreover, we analyzed whether dexamethasone might have an immediate effect on virus replication, due to its catabolic function, its inhibitory effect on pro-inflammatory pathways at the transcriptional level and inhibition of mTOR (Shah *et al*, 2000; Wang *et al*, 2006; Quatrini & Ugolini, 2021). Furthermore, we tested sorafenib, a broad-spectrum kinase inhibitor. Virus replication in presence or absence of the respective drugs was assessed by immunofluorescence staining of SARS-CoV-2 N protein at 24 hpi. While neither metformin nor dexamethasone had an inhibitory effect on virus replication, we found that sorafenib could inhibit SARS-CoV-2 replication with and IC50 concentration of 765 nM (Fig. 6D). This observation is in line with previous studies (Klann *et al*, 2020). Furthermore, GC376 also potently inhibited SARS-CoV-2 replication with an IC50 concentration of 244 nM (Fig. 6D), which is in line with recent findings by others (Vuong *et al*, 2020; Fu *et al*, 2020).

In conclusion, our 3CL^pro^ probe cleavage assay can be applied to screen for direct inhibitors of SARS-CoV-2 replication and characterize the dynamics of inhibitor binding and release. We found that direct as well as indirect interference with SARS-CoV-2 replication was effective in our experimental system. Interestingly, we observed a striking difference between indirect inhibitors of the mTOR pathway and the multi-kinase inhibitor sorafenib, suggesting that a specific subset of kinases could serve as optimal target(s) to counteract SARS-CoV-2 replication.

## Discussion

Our analysis of SARS-CoV-2 and host cell transcriptomes showed an initial upregulation of transcription in a fraction of genes starting between 0 and 2 hpi, rapid replication of virus transcripts up to levels comparable to the whole host cell transcriptome at 12 hpi, followed by transcriptional downregulation in a broad range of genes between 24 and 48 hpi, which coincided with cell death and virus release. To focus on the timing and sequence of the transcriptional response, we developed an approach based on fitting profile functions to transcriptomic measurements thereby extracting time points of maximal transcriptional regulation. Measurements of virus transcripts, virus proteins and information about affected cellular pathways were combined to investigate the sequence of interactions between SARS-CoV-2 and host cells.

To identify cellular processes that are essential for explaining SARS-CoV-2 replication in our experimental system, we developed a model that was devoid of mechanistic details in contrast to previous ODE-models of RNA virus replication that included aspects as positive and negative RNA strands, virus particle formation or shuttling between cellular compartments (Binder *et al*, 2013; Aunins *et al*, 2018; Zitzmann *et al*, 2020; Lopacinski *et al*, 2021). Modeling showed that taking into account the inhibition of the transcription of anti-viral response genes by virus proteins was required to explain our experimental dataset. Modeling further suggested that the cellular anti-viral response resulted in a decrease of virus transcripts, but was nevertheless insufficient in preventing virus production and cell death. Our analysis and modeling inferred a sequence of events comprising (1) rapid upregulation of anti-viral response genes, (2) accumulation of virus proteins, and (3) the expression of anti-viral response genes depending on virus proteins. This indicates that delaying the replication of virus transcripts or synthesis of virus proteins by interfering with virus replication could serve as a strategy to sustain the anti-viral response and increase its efficiency. Additionally, modeling suggested that the largest part of virus transcripts is not required for synthesis of virus proteins. This leads to the naïve hypothesis that the largest part of virus RNA might have the simple physiological purpose as danger-associated molecular pattern (DAMP) species triggering an inflammatory response after cell lysis, thereby supporting viral spread through tissue damage.

Focusing on the dynamics of the transcriptional response in cells infected with SARS-CoV-2 and the temporal order of affected cellular processes, we could observe a rapid sequence of upregulation in pathways related to cellular inflammation shortly after virus entry. This was followed by upregulation in processes associated with protein synthesis and production of chemical energy, all of which occurred before virus transcript levels strongly increased. Upregulation in the latter processes likely results from a particularly high energy demand during virus replication. Hijacking of the cellular metabolism related to processes such as glycolysis, nucleotide or lipid synthesis was observed before in several viruses and can be compared to metabolic reprogramming in cancer cells (Thaker *et al*, 2019). Host pathway upregulation mostly occurred prior to the time when large virus transcript levels were present and decreased after 24 hpi when cells died and virus particles were released. Besides an early transcriptional upregulation in pathways related to inflammation, such as the IL-17 or TNF signaling pathways, we observed an early transcriptional upregulation in pathways depending on kinase signaling such as the PI3K/Akt/mTOR or MAPK pathways. A causal link between the early transcriptional upregulation in the PI3K/Akt/mTOR pathway and later upregulation of mitochondrial and ribosomal genes can be hypothesized because it is known that this pathway is involved in the control of the biogenesis of mitochondria and ribosomes (Iadevaia *et al*, 2012). Previously, it was shown that NF-κB signaling is linked to upregulation of mitochondrial respiration (Mauro *et al*, 2011). It was further observed that PI3K/mTOR inhibition decreased c-Myc induction and inhibited influenza virus replication (Smallwood *et al*, 2017), which indicates that transcriptional upregulation of this pathway is part of the host cell responses that support virus replication. A recent study showed that SARS-CoV-2 limits AMPK/mTORC1 activation and autophagy, and that virus propagation is inhibited by the Akt inhibitor MK-2206 (Gassen *et al*, 2020). Taken together, the host cell response dynamics is characterized by early activation of pathways involved in inflammation and kinase signaling followed by transcriptional upregulation of processes apparently supporting virus replication.

Interestingly, we observed downregulation of xenobiotic metabolism involving cytochrome P450 enzymes in response to SARS-CoV-2 infection. It was previously observed that exposure of cells to inflammatory cytokines such as IL-6 decreases the expression of cytochrome P450 enzymes in several tissues (Bertilsson *et al*, 2001; Aitken & Morgan, 2007; Li *et al*, 2014; Mimura *et al*, 2015). So far, it was hypothesized but not experimentally shown that COVID-19 affects cytochrome P450 enzyme expression and drug metabolism (El-Ghiaty *et al*, 2020; Deb & Arrighi, 2021). In line with this hypothesis, we here provide experimental evidence that SARS-CoV-2 infection indeed directly affects pathways related to the bioavailability and metabolism of drugs.

To expedite the identification and characterization of direct inhibitors of virus protein maturation, we developed a cleavage probe assay that can indicate the activity of the SARS-CoV-2 3CL^pro^ and evaluated the assay using the protease inhibitor GC376. In the future, this assay could be used to screen for further 3CL^pro^ inhibiting drugs. Moreover, in our experimental setup of highly infectable Caco-2 cells, we could indeed confirm the effect of GC376 and sorafenib as direct and indirect inhibitors of SARS-CoV-2 replication, respectively (Vuong *et al*, 2020; Fu *et al*, 2020; Klann *et al*, 2020). On the other hand, metformin and dexamethasone, which are indirect inhibitors of mTOR signaling, did not prevent virus replication. This suggests that the therapeutic effect of dexamethasone is restricted to its established function in limiting the inflammatory response in COVID-19 (Tomazini *et al*, 2020; RECOVERY Collaborative Group *et al*, 2021).

Collectively, by integrating information on the dynamics of SARS-CoV-2 replication and transcription in host cells, our study sheds light on the dynamics of cellular regulatory processes involved in SARS-CoV-2 infection at a systems level. In particular, the sequential transcriptional activation of inflammatory pathways linked to the anti-viral response followed by upregulation of processes supporting virus replication and late downregulation of metabolic processes before cell death and virus release could be characterized. Our study provides additional insights into the orchestrated SARS-CoV-2 host interactions and could facilitate the development of strategies for combined drug interference with virus replication and cellular processes supporting virus biogenesis.

## Methods

### Cell culture

Caco-2 (human colorectal adenocarcinoma) cells and Vero E6 (African green monkey kidney epithelial) cells were obtained from ATCC. Cells have been tested negative for mycoplasma infection (MycoAlert Plus; Lonza, Basel, Switzerland). Caco-2, Vero E6 and HEK293T cells were maintained in Dulbecco’s modified Eagle’s medium (Invitrogen, Carlsbad, CA, USA) containing 10% fetal calf serum (Biochrom, Berlin, Germany), penicillin and streptomycin (100μg/ml; Invitrogen). HEK293T cells were transfected with X-tremeGENE HP (Roche Diagnostics, Rotkreuz, Switzerland) following the manufacturer’s instructions.

### Reagents

We used antibodies to detect the SARS-CoV-2 nucleoprotein (mouse monoclonal; Sino Biologicals, Hong Kong, China) and GFP (mouse monoclonal; Roche Diagnostics). For microscopy, the secondary antibody Alexa Fluor goat anti-mouse 568 (Thermo Fisher Scientific, Waltham, MA, USA) was applied. A Horseradish peroxidase-conjugated anti-mouse antibody (Southern Biotech, Birmingham, AL, USA) was used for immunoblotting.

### Virus preparation

SARS-CoV-2 (strain BavPat1) was obtained from the European Virology Archive. The virus was amplified in Vero E6 cells and collected at passage 3. Virus titers were determined by TCID50 assay. Caco-2 cells were infected using an MOI of 5 virus particles per cell. Medium was removed from Caco-2 cells and virus was added to cells for 1 hour at 37°C. Virus was removed, cells were washed once with PBS, and medium containing a tested inhibitor or empty medium was added to the cells.

### Determining TCID50 in Vero cells

Vero E6 cells were seeded into a 96-well plate 24 h prior to infection using 20,000 cells per well. A volume of 100 μl of viral supernatant was added to the first well, followed by seven 1:10 dilutions that were added to subsequent wells. All experiments were performed in triplicates. Infections were allowed to proceed for 24 h. At 24 hpi, cells were fixed in 2% paraformaldehyde (PFA) for 20 minutes at room temperature (RT). PFA was removed and cells were washed twice in 1×PBS and then permeabilized for 10 min at RT in 0.5% Triton X-100. Cells were blocked in a 1:2 dilution of LI-COR blocking buffer (LI-COR, Lincoln, NE, USA) for 30 min at RT. Cells were stained with 1/1,000 dilution anti-dsRNA (J2) for 1 h at RT. Cells were washed three times with 0.1% Tween in PBS. Secondary antibody (anti-mouse CW800) and DNA dye Draq5 (Abcam, Cambridge, UK) were diluted 1/10,000 in blocking buffer and incubated for 1 h at RT. Cells were washed three times with 0.1% Tween/PBS. Washing buffer was replaced by 1×PBS (without Tween), and samples were imaged using a LI-COR imager.

### RNA extraction, viral RNA quantification and sequencing

RNA was extracted from infected or mock-treated Caco-2 cells at 0, 1, 2, 4, 7, 12, 24 and 48 hpi using the Qiagen RNAeasy plus extraction kit (Qiagen, Hilden, Germany). For quantifying SARS-CoV-2 genome abundance in mock samples, cDNA was made using iSCRIPT reverse transcriptase (BioRad, Hercules, CA, USA). q-RT-PCR was performed using iTaq SYBR green (BioRad) as per manufacturer’s instructions, TBP was used as housekeeping gene and for normalization (COV1 primers: for 5’-GCCTCTTCTGTTCCTCATCAC-3’, rev 5’-AGACAGCATCACCGCCATTG-3’; TBP primers: for 5’-CCACTCACAGACTCTCACAAC-3’, rev 5’-CCACTCACAGACTCTCACAAC-3’; Eurofins, Luxemburg). RNA samples were stored at −80°C. DNA libraries were prepared at GeneWiz Inc. (Leipzig, Germany) using the NEBnext Ultra II RNA directional Kit (New England Biolabs GmbH, Frankfurt, Germany). Paired-end sequencing of 2×150bp was performed at GeneWiz Inc. using an Illumina NovaSeq 6000 instrument (Illumina, San Diego CA, USA).

### Microscopy

To analyze SARS-CoV-2 N protein expression in Caco-2 cells by immunofluorescence, microscopic images were taken at 0, 4, 24 and 48 hpi using a Nikon Eclipse Ti-S fluorescence microscope (Nikon, Tokio, Japan). Viable and dead cells were distinguished and quantified based on DAPI staining. Live-cell experiments were performed on a Nikon Ti inverted microscope, equipped with a CSU-22 Yokogawa confocal spinning disc slider (Yokogawa Electric Corporation, Tokyo, Japan), a 60× Plan Apo NA 1.4 objective lens (Nikon), a Hamamatsu C9100-02 EMCCD camera (Hamamatsu Photonics, Hamamatsu, Japan), and the Volocity software (PerkinElmer; Waltham, MA, USA). Fluorescence of mGFP was excited at 488 nm and emission was collected through a 527/55 emission filter (Chroma Technology Corp, Bellows Falls, VT, USA), and mCherry fluorescence was excited at 561 nm and emission was collected with a 615/70 filter (Chroma Technology Corp, Bellow Falls, VT, USA). For time-resolved experiments, images were recorded at a time interval of 5 minutes. Microscopic images were evaluated with ImageJ software (NIH, Bethesda, MA, USA).

### Cloning of 3CL^pro^ expression vectors and cleavage probes

Cleavage probes for monitoring the SARS-CoV-2 main protease activity were created by inserting 3CL^pro^ targeted sequences in between an NES sequence (MNLVDLQKKLEELELDEQQ) and mGFP. To this end, the NES-mGFP encoding vector previously reported by us (Beaudouin *et al*, 2013) was linearized via unique *Age*I/*Not*I restriction sites, and cleavage probes were inserted by oligo cloning. To generate a construct for co-expressing mCherry and SARS-CoV-2 main protease (3CL^pro^), a fragment encoding 3CL^pro^ was first obtained as a double-stranded DNA fragment (gBlock; IDT, San José, CA, USA) and cloned into pcDNA3.1(−) via unique *Nhe*I/*Not*I restriction sites. A fragment encoding mCherry followed by a P2A peptide sequence was also obtained as gBlock. The 3CL^pro^ encoding vector was then linearized 5’ of the 3CL^pro^ start codon via *Nhe*I and the mCherry-P2A fragment was inserted via Gibson Assembly, hence yielding a vector encoding mCherry-P2A-3CL^pro^.

### Co-expression of 3CL^pro^ with cleavage probes and immunoblotting

One day before transfection, 10^5^ cells per well were seeded in 6-well plates. Cells were co-transfected with constructs encoding a cleavage probe and either (1) 3CL^pro^-2A-mCherry (2) a control vector lacking 3CL^pro^ (Kallenberger *et al*, 2014). For harvesting lysates, two days after transfection, plates were transferred on an ice-cold metal block, washed in ice-cold 1× PBS before treatment with ice-cold lysis buffer [20 mM tris-HCl (pH 7.5),150 mM NaCl, 1 mM phenylmethylsulfonyl fluoride (Millipore Sigma, St. Louis, MO, USA), protease inhibitor cocktail (Roche Diagnostics), 1% Triton X-100, and 10% glycerol], and harvested with cell scrapers (BD Biosciences, Franklin Lakes, NJ, USA). Lysates were analyzed with SDS – polyacrylamide gel electrophoresis gels (Invitrogen). Proteins were transferred onto polyvinylidene difluoride (PVDF) membranes (Millipore Sigma) by wet blotting. A primary antibody recognizing GFP (Roche Diagnostics) and a horseradish peroxidase-conjugated secondary antibody (Southern Biotech) were used to probe membranes. Detection was performed using the Pico Chemiluminescent Substrate from Thermo Scientific and a charge-coupled device camera (Intas, Göttingen, Germany).

### Bioinformatics and statistical analysis

Read pairs from triplicate samples of infected cells at 0, 1, 2, 4, 7, 12, 24 and 48 hpi and additional mock samples at 4, 12 and 48 hpi were mapped to a merged reference comprising the human reference genome (GRCh38.p13, NCBI build 38 path release 13 obtained from Genome Reference Consortium) and SARS-CoV-2 reference (NC_045512.2, NCBI reference sequence for SARS-CoV-2 isolate Wuhan-Hu-1). For read alignment, the STAR software was used with default settings (Dobin *et al*, 2013). Subsequently, BAM files were split by their reference using the SAMtools software suite (Li *et al*, 2009) and counted separately using the featureCounts function from the Subread package (Liao *et al*, 2019). Host reads were counted by the ‘Exon’ feature, whereas SARS-CoV-2 mapping reads were counted by the ‘CDS’ feature. Transcript read counts were processed using ‘DESeq2’ (Love *et al*, 2014). At first, log2 fold changes were calculated relative to a reference comprising measurements of samples at 0 and 1 hours. Then, for background correction of time series, log2 fold changes of measurements from uninfected samples were subtracted from log2 fold changes of measurements from infected samples (Supplementary Fig. S9). To this end, mock values were linearly interpolated at 1, 2, and 7 hpi. To estimate errors of background-corrected values *x* at all time points, standard errors of gene expression data from mock samples were fitted by a linear error model *ε*(*x*) = *m*_1_*x* + *m*_2_*max*(*x*). The parameters *m*_1_ and *m*_2_ were estimated for each gene by performing 100 multi-start local optimizations. Calibrated error models were used to estimate errors of interpolated mock values at 1, 2 and 7 hpi. Finally, errors of background-corrected values were obtained from error propagation using standard errors of measurements from infected samples and mock samples. Background-corrected and interpolated log2 fold changes were then used for GO term enrichment analysis (Ashburner *et al*, 2000; Gene Ontology Consortium, 2021). For cluster analysis, and for visualizing transitions between GO-terms, log2 fold changes were interpolated in hourly time intervals between 0 and 48 hpi. Overrepresented GO terms were inferred using the ‘goana’ and ‘topGO’ function from the limma package (Ritchie *et al*, 2015). As input, we used all genes with an absolute log2 fold change larger than one. Besides the gene label, we explicitly used the trend or abundance parameter in the ‘goana’ function (mean expression of gene *i* at timepoint *j*). We subsequently selected the top GO terms for each discrete time point and visualized negative log-scaled p-values in a clustered heatmap. We applied hierarchical clustering using the Euclidean distance norm and complete linkage as implemented in the ‘hclust’ function of the ‘stats’ package.

### Analyzing the dynamics of transcription changes

To extract time points when gene expression was strongly affected and determine amplitudes of expression changes in all expressed host cell genes, profile functions were defined. Log2 fold changes were fitted by four profile functions

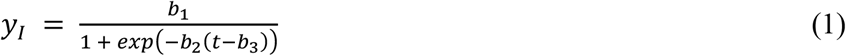

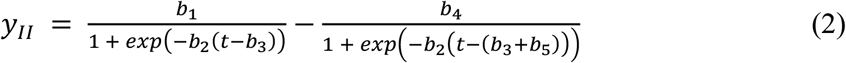

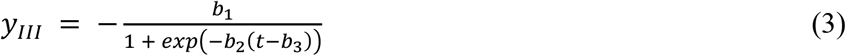

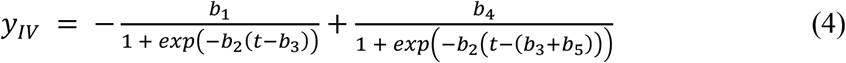

to describe continuously increasing (*y_I_*) transiently increasing (*y_II_*), continuously decreasing (*y_III_*) or transiently decreasing expression (*y_IV_*) of genes. Eqs. (1–4) were fitted to log2 fold changes of all *N* = 13,322 expressed genes. In each case, all four profile functions were fitted.

The optimal function was selected using the Bayesian information criterion

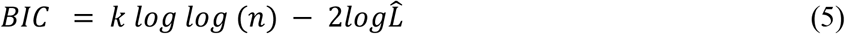

that depends on the logarithm of the likelihood function maximum 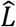, the number of parameters *k* and the number of datapoints *n*. Parameters *b_i_* were log-scaled to facilitate convergence of fits. For each profile function, 100 multi-start local optimizations were performed.

### Mathematical modeling

Model simulations were performed with custom scripts in MATLAB (The Mathworks, Natick, MA, USA). The MATLAB toolbox PottersWheel was used for model fitting (Maiwald & Timmer, 2008). Models described four species, virus transcripts *V*, virus proteins *P*, mRNAs of anti-viral genes *m_A_* and anti-viral proteins *A* (Fig. 5A). Model versions comprised between 12 and 14 estimated parameters. The model variable *V* was associated with measured virus transcript counts. The GO term ‘Defense response to virus’ (GO:0051607) was used to derive an observable for the model species *m_A_*. After fitting profile functions, as described above, to all expressed genes associated with this GO term, strongly regulated genes were selected based on log2 fold change amplitudes of *log*_2_ *f*. *c*. ≥ *log*_2_(3/2) (Supplementary Fig. S4).

No additional scaling factors were used to relate experimental measurements in arbitrary units to concentrations or molecule numbers of model species. Therefore, scaling constants implicitly entered the kinetic parameters for describing species turnover. The only non-zero valued model species was [*V*_0_], the initial level of *V* after infection of cells. In total, 18 model variants were iteratively developed (Supplementary Figs. S5–7). Each variant was calibrated by performing 5000 multi-start local optimizations. Model selection was performed based on the BIC.

The initial model version (Variant 1, Supplementary Fig. S5) described replication of *V*, translation of *V* to *P*, induction of *m_A_* translated to *A*, and inhibition of *V* synthesis by *A*. It was extended by six additional parts: (1) positive feedback of *P* on replication of *V* (Variant 2), (2) positive feedback of *P* on virus replication by virus release and influx (Variant 3), (3) negative feedback of *P* on the transcription of *m_A_* (Variant 4), (4) negative feedback of *P* by cleavage of *m_A_* (Variant 5), (5) negative feedback of *P* on the synthesis of *A* (Variant 6), or (6) dependency of the expression of *m_A_* on a threshold level of *V* (Variant 7). Model selection showed that ‘Variant 4’ was superior compared to the other variants (Supplementary Fig. S5B, C).

In a second model extension step, the remaining additional parts were added to Variant 4 (Variants 4.1–5; Supplementary Fig. S6). Additional inclusion of the negative feedback of *P* by cleavage of *m_A_* resulted in an improved fit indicated by a slightly decreased chi-square measure (Supplementary Fig. S6B). None of the additional model extensions, however, resulted in a decrease in BIC (Supplementary Fig. S6C).

In a third step, it was tested whether model Variant 4 could be simplified by more parsimonious model variants containing less parameters without decreasing fit quality. To this end, the following six model simplifications were tested: (1) description of *A* turnover by one turnover parameter instead of separate parameters for synthesis and degradation of *A* (Variant 4.0.1), (2) description of the synthesis of *A* by mass-action instead of Michaelis-Menten (MM) kinetics (Variant 4.0.2), (3) mass-action instead of MM-kinetics for describing synthesis of *P* (Variant 4.0.3), (4) mass-action kinetics for describing synthesis and degradation of *A* by single turnover parameter (Variant 4.0.4), (5) single parameter for describing turnover of *A* and mass-action instead of MM-kinetics for synthesis of *P* (Variant 4.0.5), (6) mass-action kinetics for synthesis of *A* as well as *P*. Model selection showed that ‘Variant 4.0.1’ was superior compared to the other variants and subsequently regarded as optimal model variant (Supplementary Fig. S7B, C).

Model equations of all variants are listed in Supplementary Table S1; parameter estimates of the optimal model variant ‘4.0.1’ including allowed parameter intervals and 1σ-confidence intervals estimated based on the inverse of the Hessian matrix are given in Supplementary Table S2.

## Acknowledgements

We thank Thomas Wolf and Benedikt Brors for comments and valuable discussions. This work was supported by the fightCOVID-19 initiative of the German Cancer Research Center (Heidelberg, Germany). Furthermore, this work was supported by research grants from the Bundesministerium Bildung und Forschung (BMBF): project numbers 01KI20239B to MS, 031L0270 to SK (Computational Life Sciences program) and 01KI20198A to SB, the Deutsche Forschungsgemeinschaft (DFG): project numbers 416072091 to MS, 415089553 (Heisenberg program), 240245660 (SFB1129) to SB, and 453202693 to DN, the state of Baden-Wuerttemberg (AZ: 33.7533.-6-21/5/1) to SB, and the Aventis foundation to DN.

## Author contributions

SK and DN conceived the study and refined it together with MS, SB and RE. MS performed SARS-CoV-2 infection experiments and generated samples for RNA sequencing. LA, FS and SK analyzed RNA sequencing data. SK performed mathematical modeling. DN and JM generated the 3CL^pro^ cleavage probes; SK, LA and CDP performed cleavage probe and immunoblotting experiments. RE, SB, SK and DN secured funding. SK wrote the manuscript with support by MS and DN. All authors approved the final manuscript.

## Data availability statement

Transcriptomic data, MATLAB and R scripts as well as vectors encoding cleavage probes and 3CL^pro^ are available from the corresponding authors upon reasonable request.

