## Supplementary Material for "Dynamics of SARS-CoV-2 host cell interactions inferred from transcriptome analyses"

### Co-corresponding authors

**This file includes**

Supplementary note S1

Supplementary figures S1 to S9

Supplementary tables S1 to S3

#### Supplementary note S1

We performed RNA-seq experiments in two Caco-2 lineages from two independent sources (University Hospital Heidelberg, EMBL Heidelberg). In one of the Caco-2 cell lineages, an infectivity of 96% was achieved at 24 hpi when using five virus particles per cell but only 15% in the other lineage (Supplementary Fig. S1A). Infectivity was determined based on immunostaining of SARS-CoV-2 N protein. At 24 hpi, in weakly infected Caco-2 cells, only 17% of the abundance of SARS-CoV-2 genomes was detected using q-RT-PCR (Supplementary Fig. S1B). In each of the two cell lines, cultured under identical conditions (see Methods), 9 uninfected samples were analyzed. Interestingly, the two lineages showed a 6-fold difference in ACE2 and 3-fold difference in cathepsin B expression, both of which are host cell proteases relevant to virus cell entry (Supplementary Fig. S1C, D). No differences were observed in expression of TMPRSS2, furin and cathepsin L. Hence, due to its superior infectivity, the high ACE2/cathepsin B Caco-2 line was employed in further experiments. Cell line identity of both Caco-2 cell lineages was confirmed based on single nucleotype polymorphism prototyping (Multiplexion GmbH, Heidelberg, Germany).

**Supplementary figures**

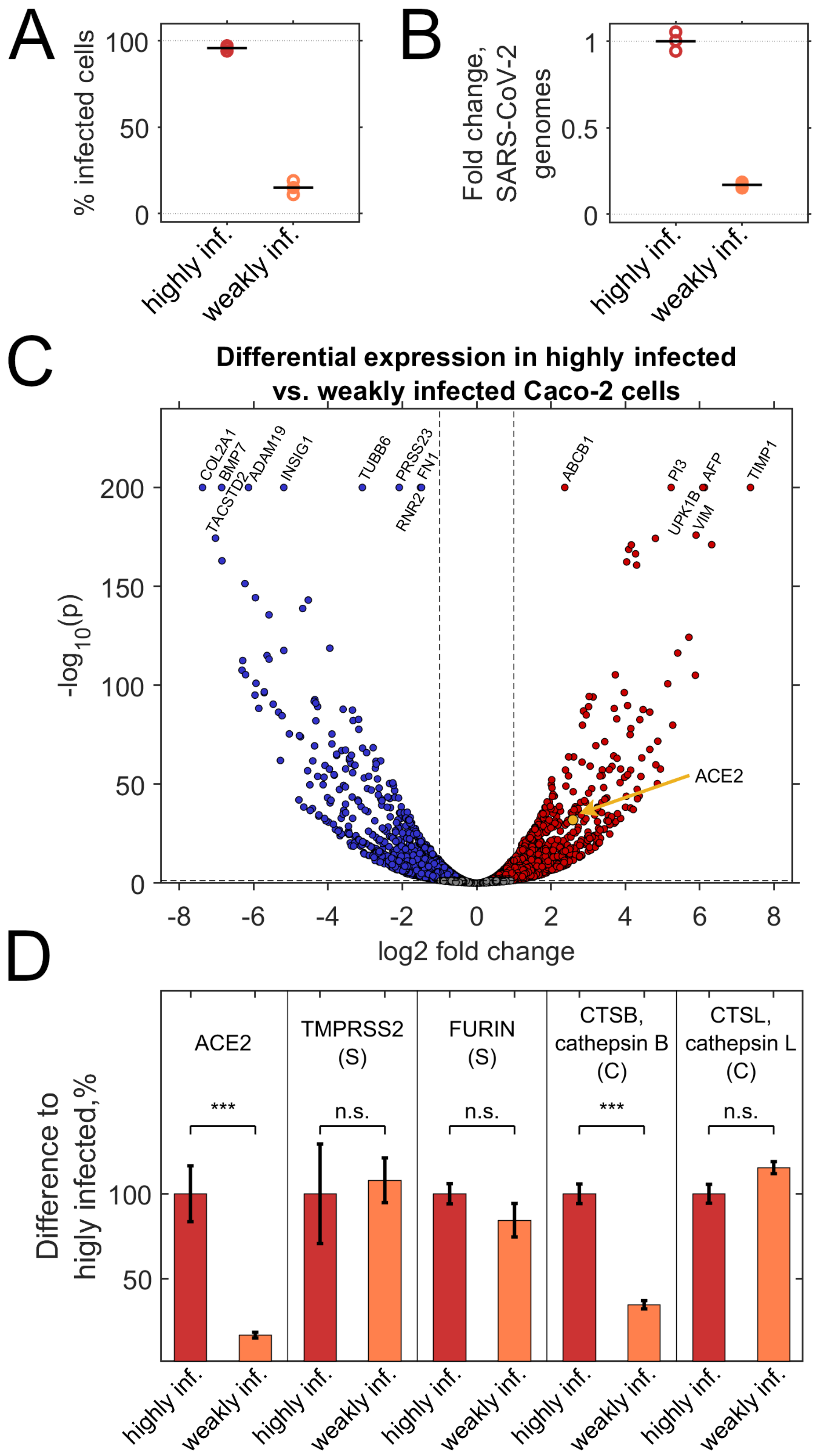

**Figure S1. Differential expression analysis of highly vs. weakly infected Caco-2 cells.** (**A**) Highly infected Caco-2 cells showed an infection level of 96%, weakly infected Caco-2 cells an infection level of 15% (Averages of n=3 replicates). Infection levels determined based on immunostaining of SARS-CoV-2 N protein. (**B**) Normalized SARS-CoV-2 genome abundance assessed by PCR. Weakly infected Caco-2 cells showed a genome abundance of about 17% of highly infected Caco-2 cells. (**C**) Log2 fold changes in gene expression and p-values of a negative binomial test followed by Benjamini-Hochberg (BH) correction for multiple testing (values were cut off for $p<10^{-200}$). Names of top differentially expressed genes are indicated. Notably, ACE2 expression was elevated in highly infected cells (indicated by arrow). We speculate that the increased expression of molecular markers as AFP, TIMP1 and VIM in highly infected, relative to weakly infected Caco-2 cells, might be indicative of cellular selection related to cancer progression in the Caco-2 cell line. (**B**) Expression of ACE2 and protease enzymes involved in SARS-CoV-2 infection (S, serine proteases; C, cysteine proteases). In highly infected cells, expression of ACE2 was 6-fold higher and cathepsin B was 3-fold higher than in weakly infected Caco-2 cells (***, p<0.001, negative binominal test followed by BH-correction; inf., infected; n.s., not significant).

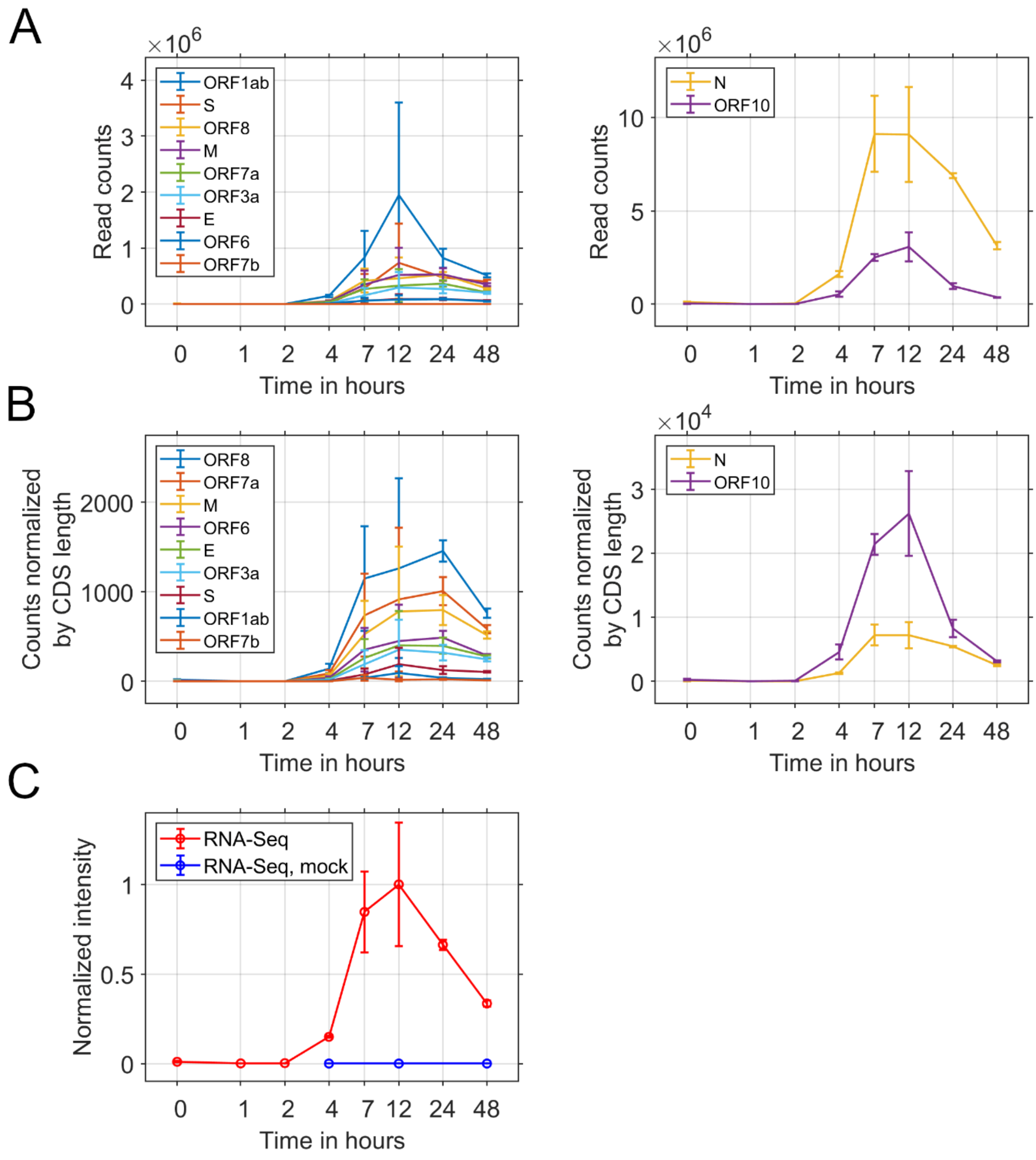


**Figure S2. Expression of SARS-CoV-2 transcripts.** (**A**) Read counts of virus transcripts (means of n=3 replicates; error bars, SEM). Read counts of most abundant transcripts N and ORF10 are shown on a separate scale (right). (**B**) Read counts normalized by lengths of coding sequences (CDS) as in (A). (**C**) Normalized average virus transcript reads (means of n=3 replicates; error bars, SEM).

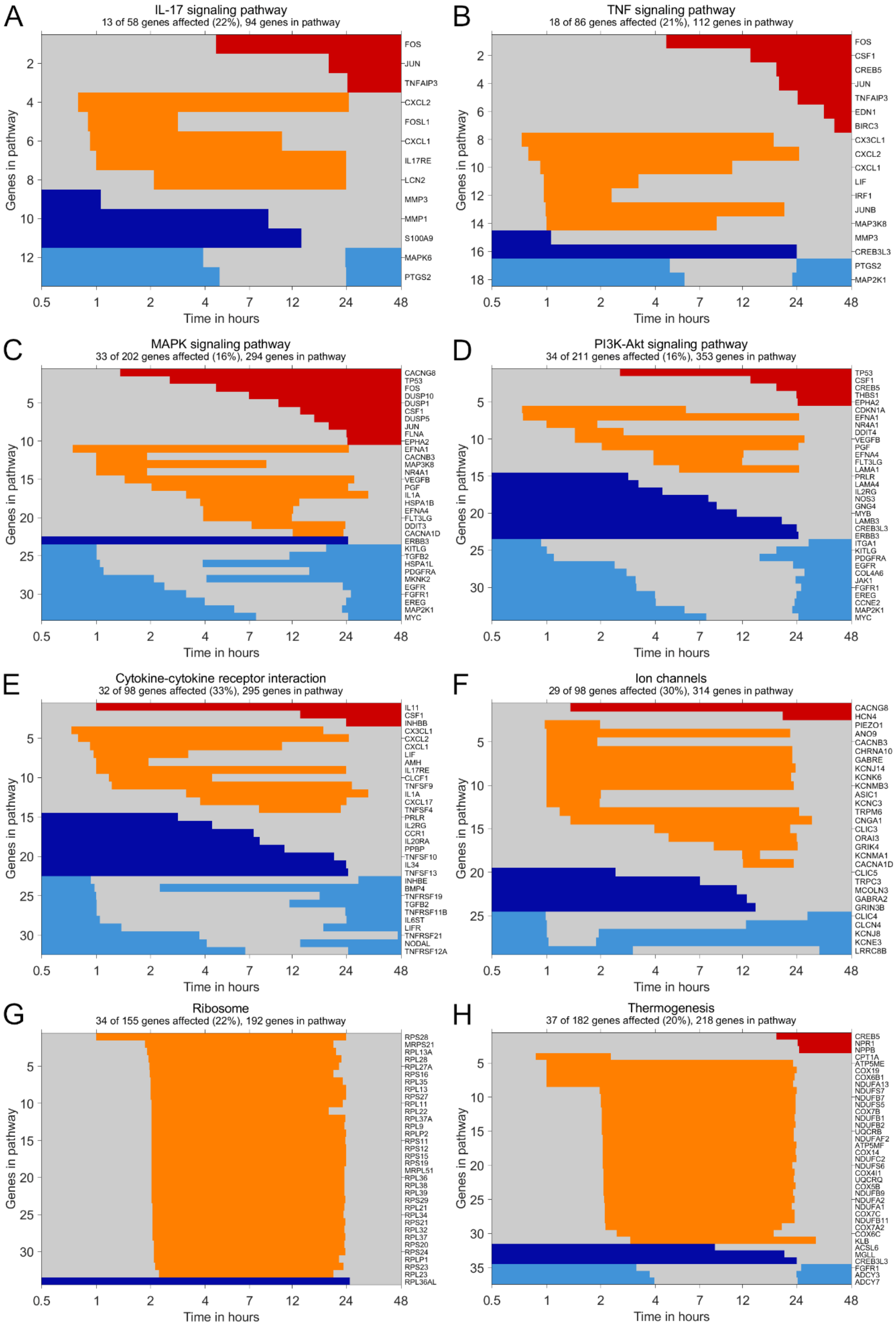

A
IL-17 signaling pathway
13 of 58 genes affected (22%), 94 genes in pathway
FOS
JUN
TNFAIP3
CXCL2
FOSL1
CXCL1
IL17RE
LCN2
MMP3
MMP1
S100A9
MAPK6
PTGS2
B
TNF signaling pathway
18 of 86 genes affected (21%), 112 genes in pathway
FOS
CSF1
CREB5
JUN
TNFAIP3
EDN1
BIRC3
CX3CL1
CXCL2
CXCL1
LIF
IRF1
JUNB
MAP3K8
MMP3
CREB3L3
PTGS2
MAP2K1
C
MAPK signaling pathway
33 of 202 genes affected (16%), 294 genes in pathway
CACNG8
TP53
FOS
DUSP10
DUSP1
CSF1
DUSP5
JUN
FLNA
EPHA2
EFNA1
CACNB3
MAP3K8
NR4A1
VEGFB
PGF
IL1A
HSPA1B
EFNA4
FLT3LG
DDIT3
CACNA1D
ERBB3
KITLG
TGFB2
HSPA1L
PDGFRA
MKNK2
EGFR
FGFR1
EREG
MAP2K1
MYC
D
PI3K-Akt signaling pathway
34 of 211 genes affected (16%), 353 genes in pathway
TP53
CSF1
CREB5
THBS1
EPHA2
CDKN1A
EFNA1
NR4A1
DDIT4
VEGFB
PGF
EFNA4
FLT3LG
LAMA1
PRLR
LAMA4
IL2RG
NOS3
GNG4
MYB
LAMB3
CREB3L3
ERBB3
ITGA1
KITLG
PDGFRA
EGFR
COL4A6
JAK1
FGFR1
EREG
CCNE2
MAP2K1
MYC
E
Cytokine-cytokine receptor interaction
32 of 98 genes affected (33%), 295 genes in pathway
IL11
CSF1
INHBB
CX3CL1
CXCL2
CXCL1
LIF
AMH
IL17RE
CLCF1
TNFSF9
IL1A
CXCL17
TNFSF4
PRLR
IL2RG
CCR1
IL20RA
PPBP
TNFSF10
IL34
TNFSF13
INHBE
BMP4
TNFRSF19
TGFB2
TNFRSF11B
IL6ST
LIFR
TNFRSF21
NODAL
TNFRSF12A
F
Ion channels
29 of 98 genes affected (30%), 314 genes in pathway
CACNG8
HCN4
PIEZO1
ANO9
CACNB3
CHRNA10
GABRE
KCNJ14
KCNK6
KCNMB3
ASIC1
KCNC3
TRPM6
CNGA1
CLIC3
ORAI3
GRIK4
KCNMA1
CACNA1D
CLIC5
TRPC3
MCOLN3
GABRA2
GRIN3B
CLIC4
CLCN4
KCNJ8
KCNE3
LRRC8B
G
Ribosome
34 of 155 genes affected (22%), 192 genes in pathway
RPS28
MRPS21
RPL13A
RPL28
RPL27A
RPS16
RPL35
RPL13
RPS27
RPL11
RPL22
RPL37A
RPL9
RPLP2
RPS11
RPS12
RPS15
RPS19
MRPL51
RPL36
RPL38
RPL39
RPS29
RPL21
RPL34
RPS21
RPL32
RPL37
RPS20
RPS24
RPLP1
RPS23
RPL23
RPL36AL
H
Thermogenesis
37 of 182 genes affected (20%), 218 genes in pathway
CREB5
NPR1
NPPB
CPT1A
ATP5ME
COX19
COX6B1
NDUFA13
NDUFS7
NDUFB7
NDUFS5
COX7B
NDUFB1
NDUFB2
UQCRB
NDUFAF2
ATP5MF
COX14
NDUFC2
NDUFS6
COX4I1
UQCRQ
COX5B
NDUFB9
NDUFA2
NDUFA1
COX7C
NDUFB11
COX7A2
COX6C
KLB
ACSL6
MGLL
CREB3L3
FGFR1
ADCY3
ADCY7
Genes in pathway
Time in hours
0.5 1 2 4 7 12 24 48

**Figure S3. Activation maps of pathways regulated upon SARS-CoV-2 infection.** Activation maps indicate time intervals of increased gene expression defined by fitting profile functions (grey, decreased level of expression; red, continuous increase; yellow, transient increase; dark blue, continuous decrease; light blue, transient decrease) as illustrated in Fig. 3. For strongly regulated genes (absolute value of log2 f.c.≥1), limits of indicated time intervals were defined by time points of half maximal increase or decrease. (**A**–**H**) Transcription dynamics in KEGG pathways with large fractions of genes showing elevated expression. For each pathway, numbers of strongly regulated genes, expressed genes (detected by RNA sequencing) and total genes are indicated. (**A**–**D**) Transcriptional dynamics in pathways that are involved in inflammation and depend on kinase activity. (**I**–**P**) Transcription dynamics in KEGG pathways with large fractions of genes showing decreased expression. (**Q**–**U**) Transcription dynamics in pathways with large absolute numbers of strongly regulated genes: (**Q**) Coronavirus disease – COVID-19, (**R**) Peptidases and inhibitors, (**S**) Membrane trafficking, (**T**) Chromosomes and associated proteins, (**U**) Exosome.

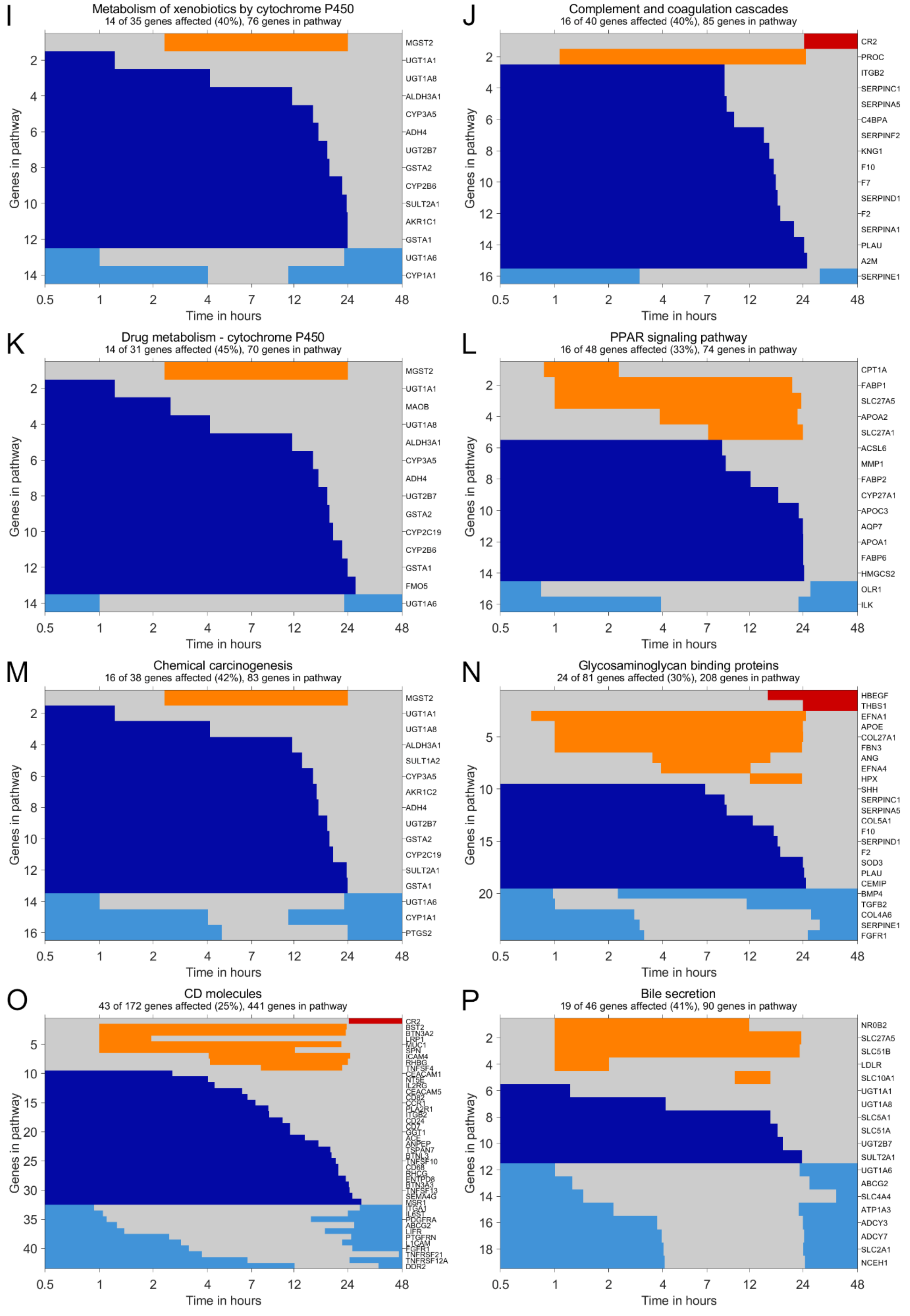


**Figure S3. (cont.)**

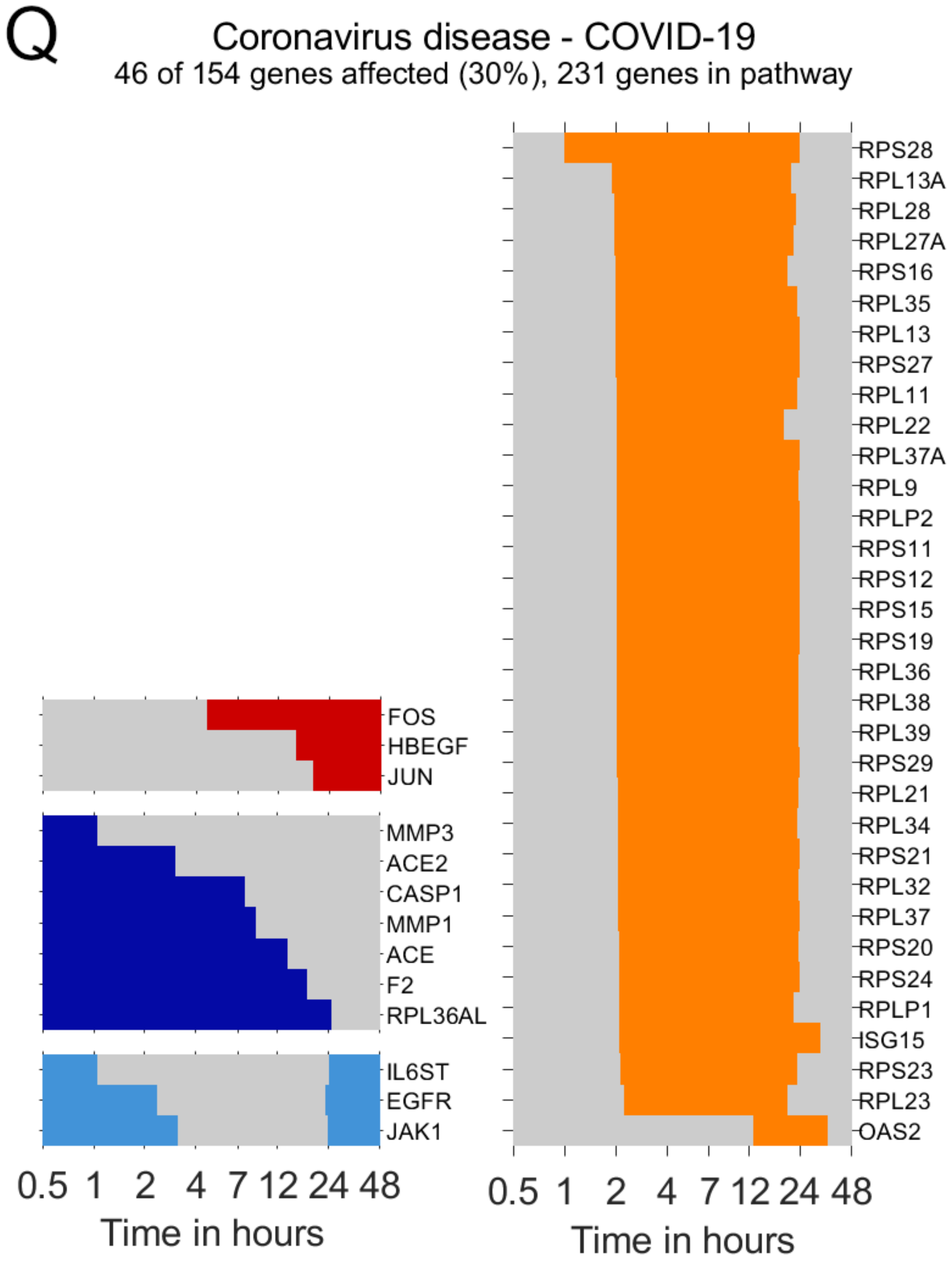


**Figure S3. (cont.)**

R

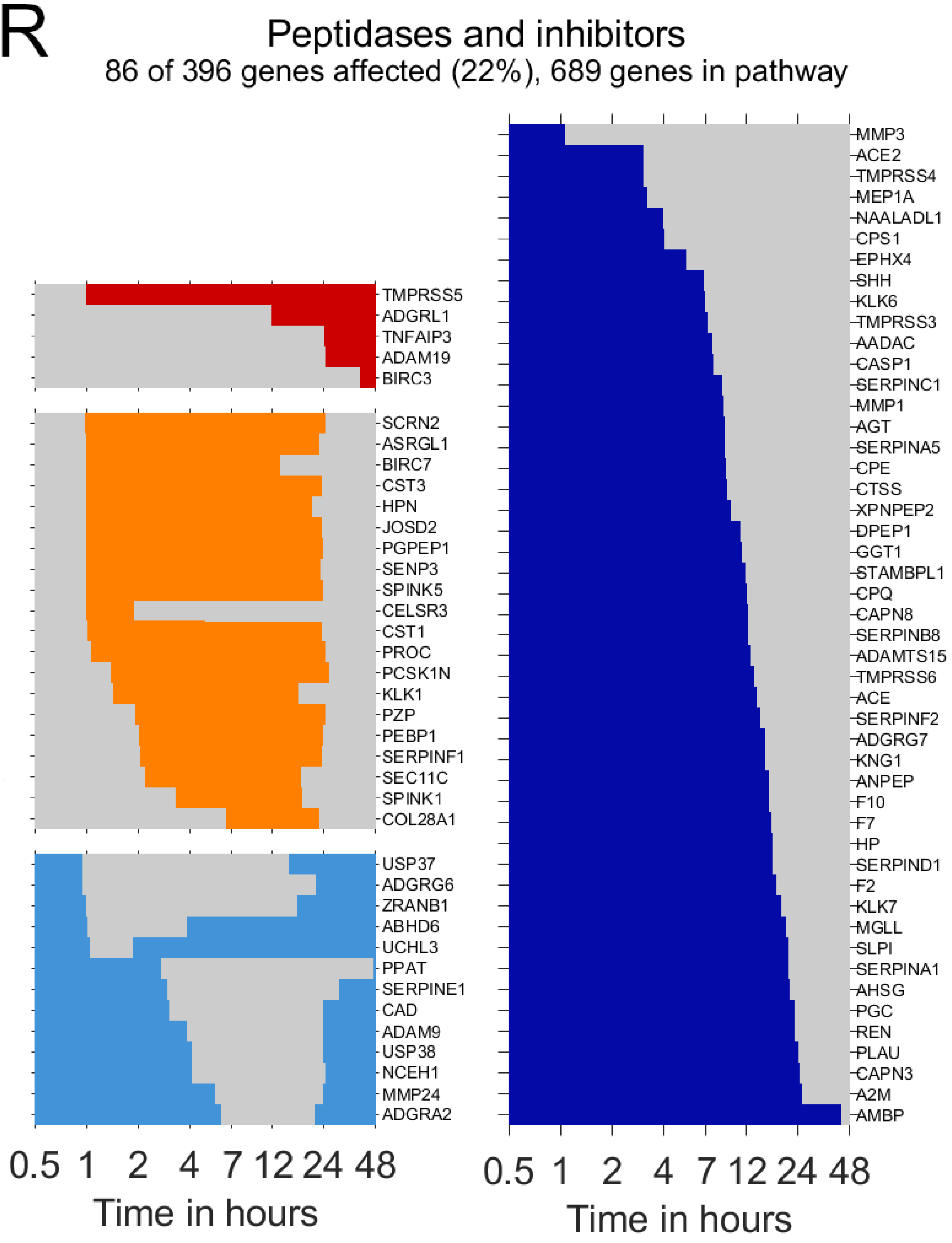


**Figure S3. (cont.)**

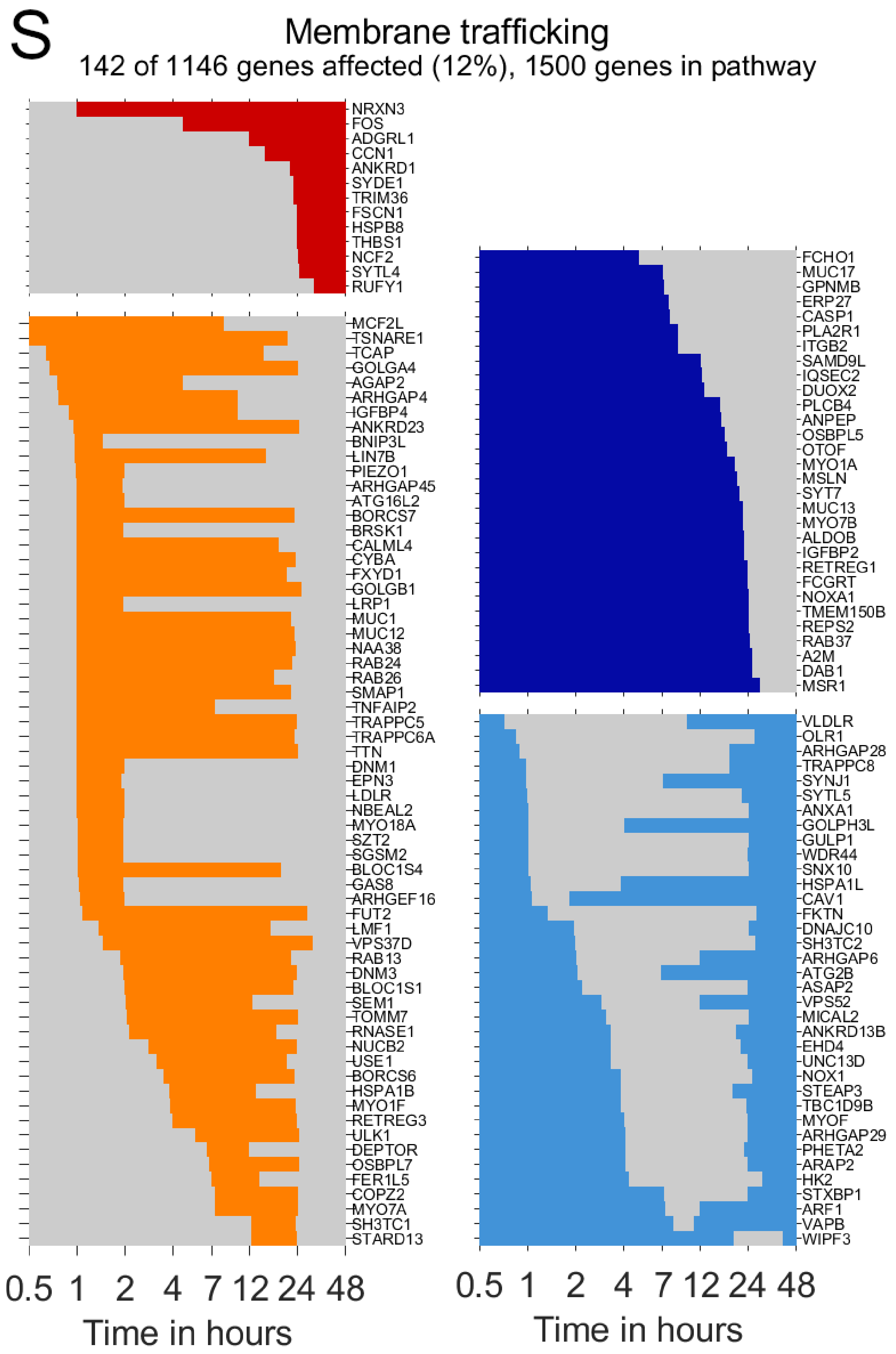


**Figure S3. (cont.)**

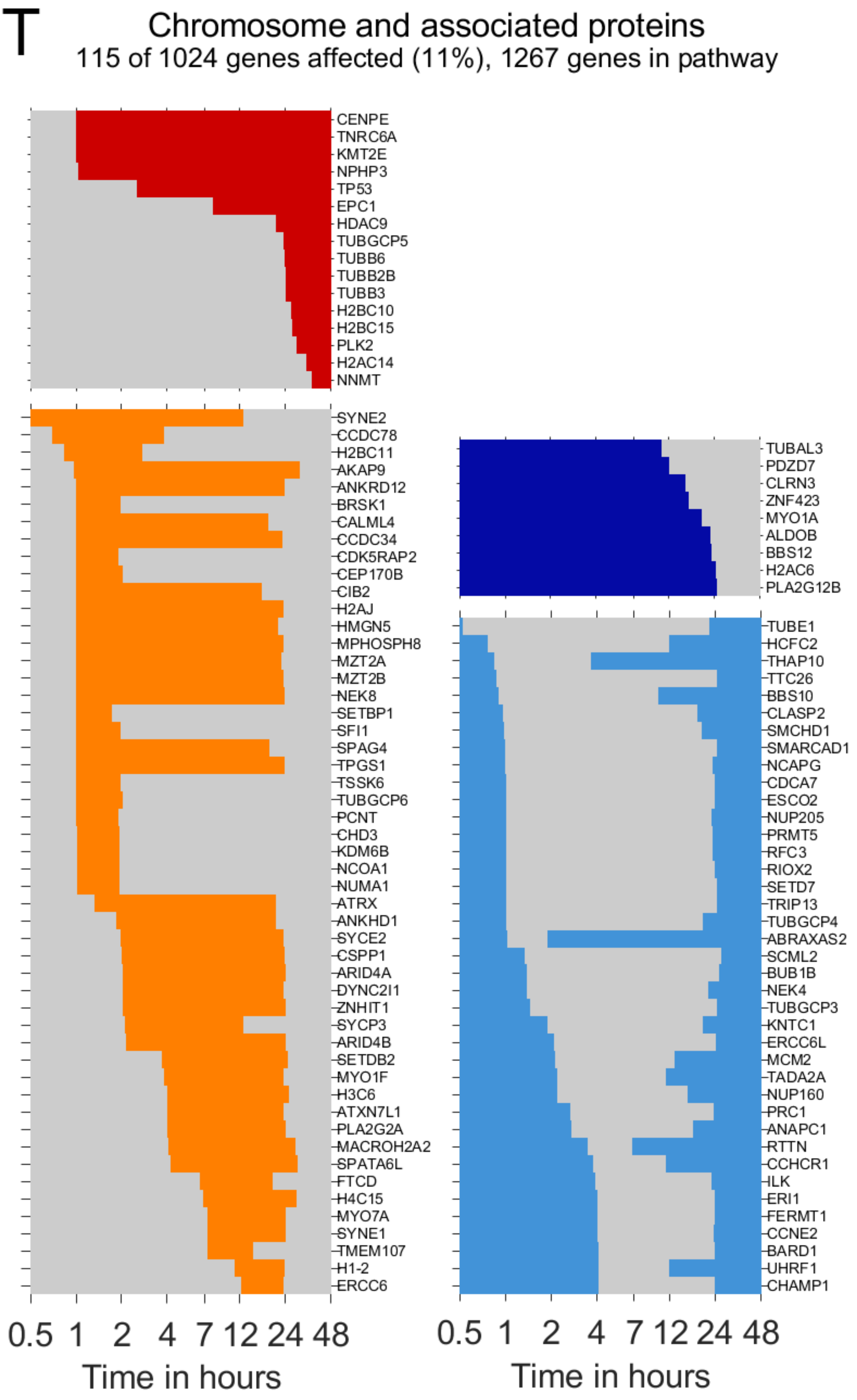


**Figure S3. (cont.)**

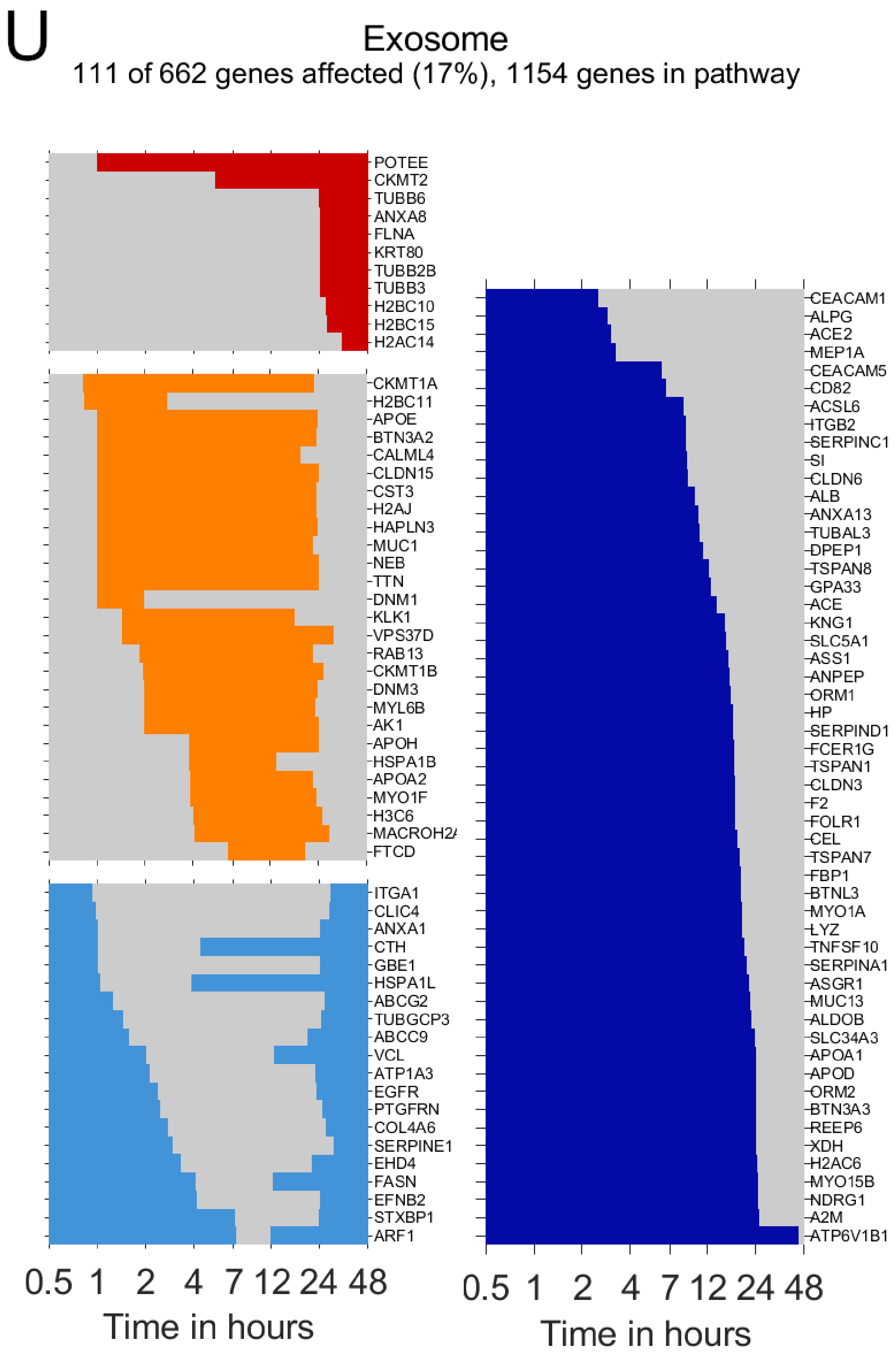


**Figure S3. (cont.)**

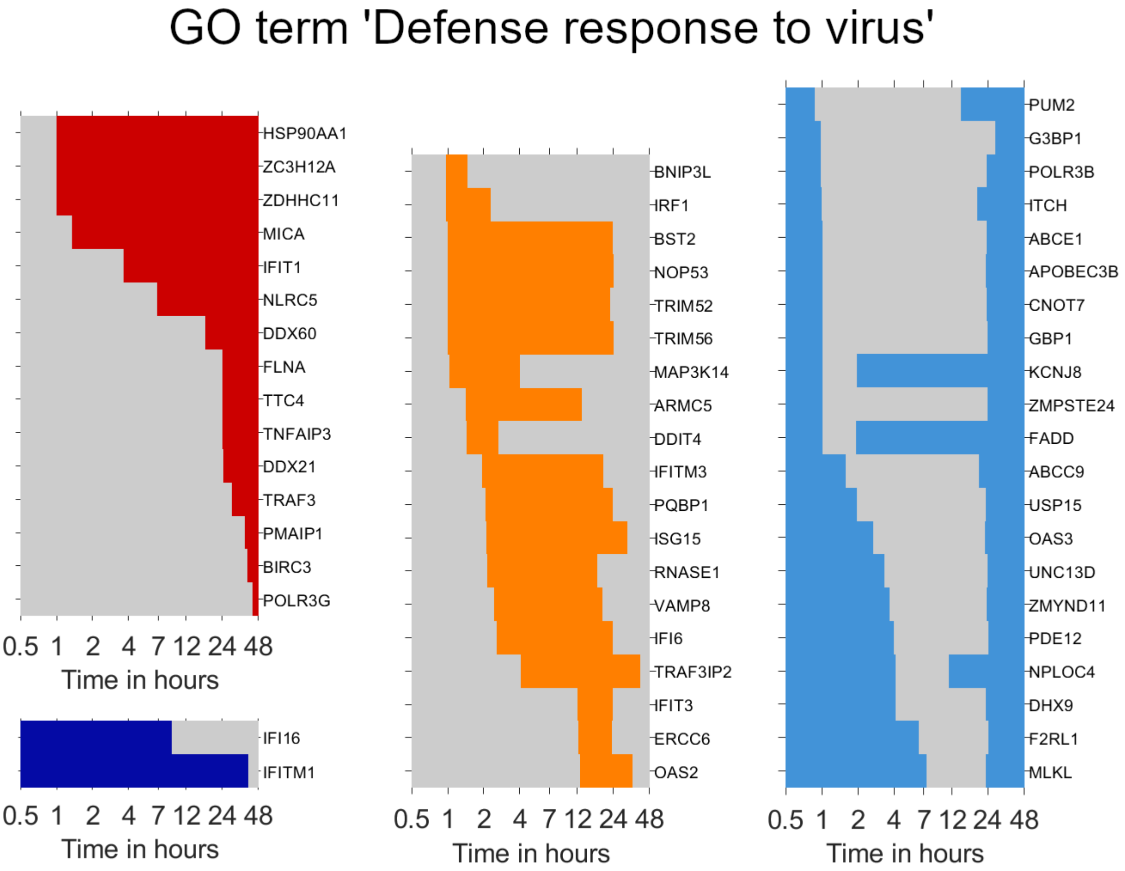


**Figure S4. Transcription dynamics in genes associated with the GO term 'Defense response to virus'.** Lines indicate time intervals of increased gene expression defined by fitting profile functions (grey, decreased level of expression; red, continuous increase; yellow, transient increase; dark blue, continuous decrease; light blue, transient decrease) as illustrated in Fig. 3. In strongly regulated genes (absolute value of $log_2\, f.c. \geq log_2(3/2)$, limits of indicated time intervals were defined by time points of half maximal increase or decrease).

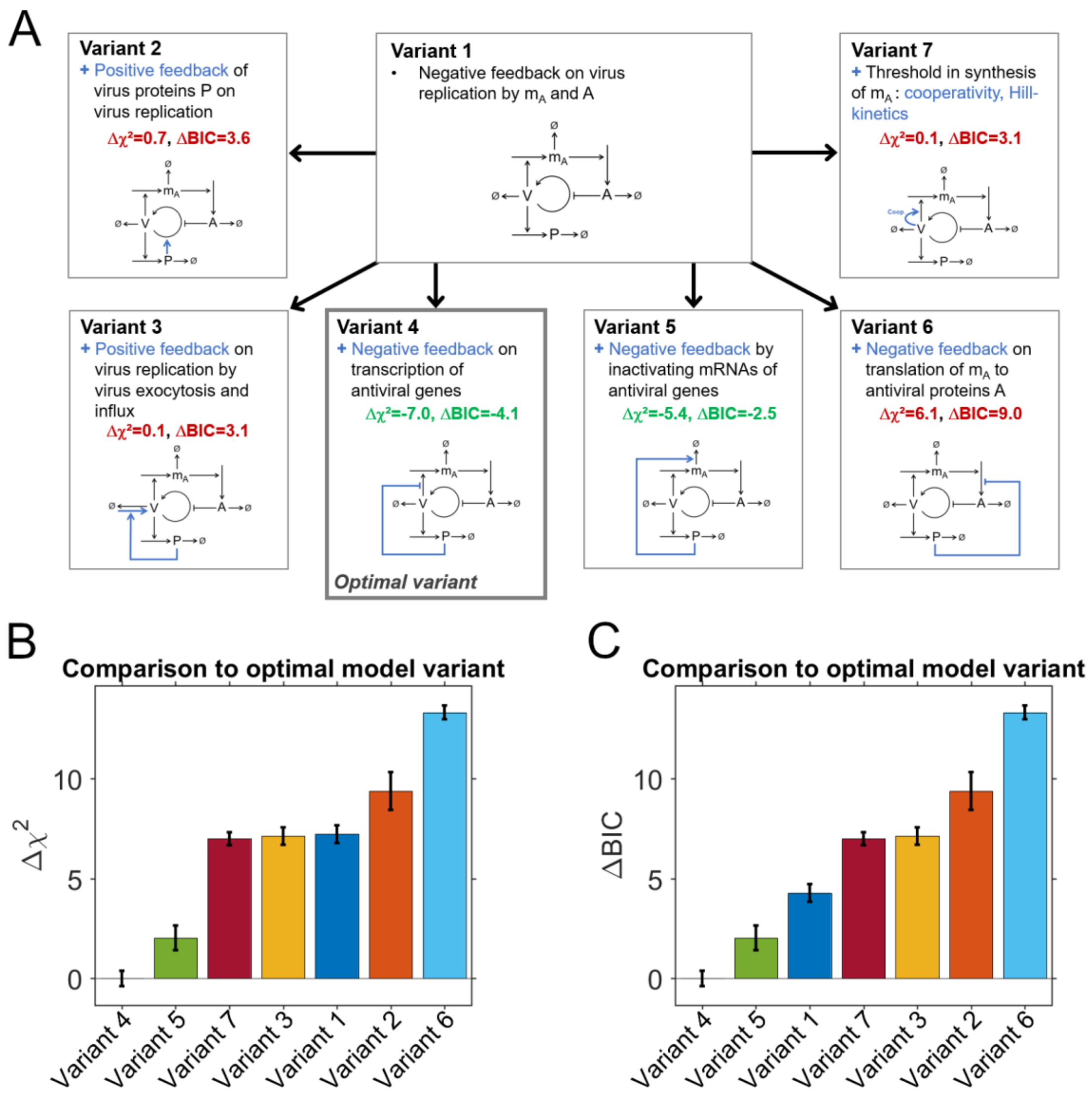


**Figure S5. First step of selecting an optimal model describing SARS-CoV-2 replication.** (**A**) The basic model ('Variant 1') describes replication of virus transcripts $V$ that induce synthesis of mRNAs of anti-viral response genes $m_A$ resulting in synthesis of anti-viral proteins $A$. These anti-viral proteins inhibit virus replication. Virus transcripts are translated to virus proteins $P$. The model was extended by reactions describing positive feedback of $P$ on virus replication ('Variant 2'), positive feedback of $P$ on influx of $V$ ('Variant 3'), inhibition of $m_A$ synthesis by $P$ ('Variant 4'), influence of $P$ on the degradation of $m_A$ ('Variant 5'), inhibition of the synthesis of $A$ by $P$ ('Variant 6') or describing $m_A$ synthesis by Hill-kinetics assuming that transcription starts in case $V$ exceeds a certain threshold ('Variant 7'). Model versions were fitted to the experimental dataset for selecting a model variant that could optimally explain the dataset based on the chi-square measure of the deviation between model and data as well as the Bayesian information criterion (BIC). (**B**) Differences in chi-square measure to the optimal model Variant 4. (**C**) Differences in BIC to the optimal model Variant 4.

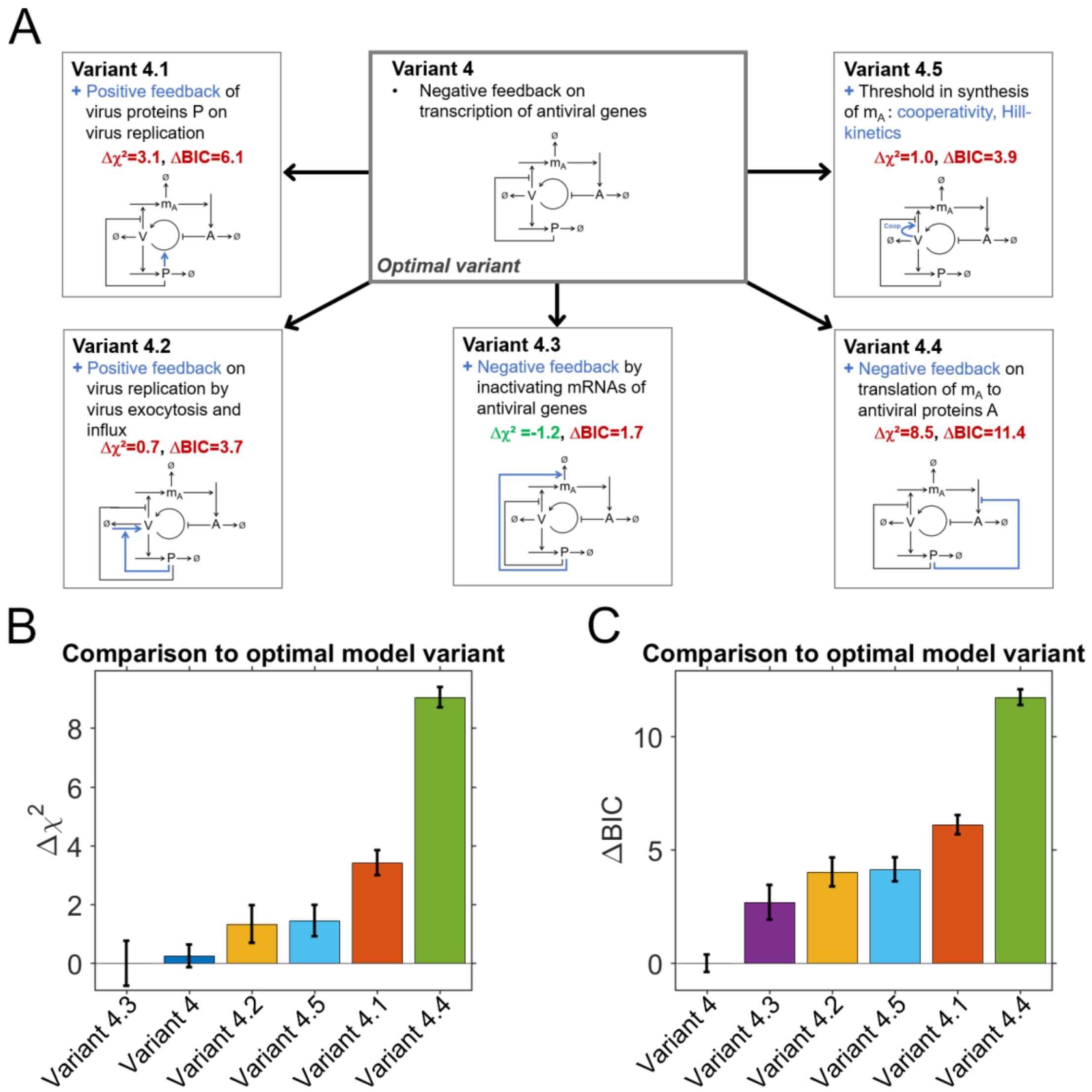


**Figure S6. Second step of selecting an optimal model describing SARS-CoV-2 replication.** (**A**) Potential further extensions of variant 4 were tested by additionally including reactions describing positive feedback from $P$ on virus replication ('Variant 4.1'), positive feedback from $P$ on virus influx ('Variant 4.2'), accelerated degradation of $m_A$ depending on $P$ ('Variant 4.3.'), inhibition of the synthesis of $A$ by $P$ ('Variant 4.4') or describing synthesis of $m_A$ dependent on $V$ by Hill-kinetics ('Variant 4.5'). Model variants were fitted to the experimental dataset. (**B**) Differences in $\chi^2$ relative to the model 'Variant 4.3' resulting in the smallest deviation between model fits and experimental data. (**C**) Differences in BIC relative to the optimal model 'Variant 4' implying that the small increase in $\chi^2$ relative to 'Variant 4.3' does not justify an extension of the model variant.

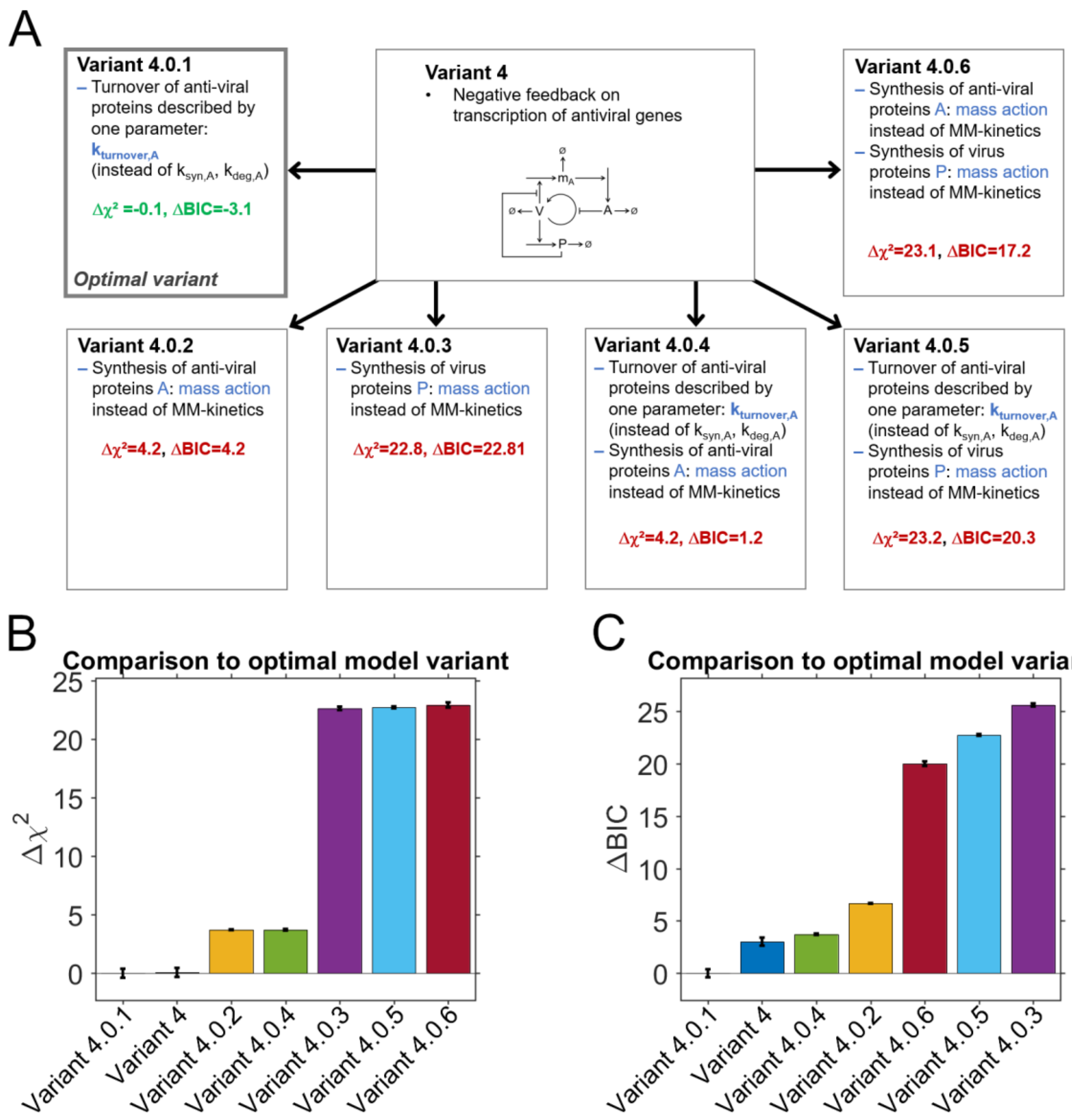


**Figure S7. Third step of selecting an optimal model describing SARS-CoV-2 replication.** (**A**) It was tested whether model 'Variant 4' could be further simplified to a more parsimonious model version with less parameters. The model was simplified by describing synthesis and degradation of $P$ ($k_{A,syn}$, $k_{A,deg}$) with just one turnover parameter ($k_{A,turnover}$, 'Variant 4.0.1'), or by replacing Michaelis-Menten (MM) kinetics by mass-action kinetics for synthesis of $A$ ('Variant 4.0.2') or $P$ ('Variant 4.0.3'). Additionally, combinations of these simplifications were tested ('Variant 4.0.4', one turnover parameter for $P$, MM-kinetics for synthesis of $A$; 'Variant 4.0.5', one turnover parameter for $P$, MM-kinetics for synthesis of $P$; 'Variant 4.0.6'; MM-kinetics for synthesis of $A$ and $P$). (**B**) Differences in chi-square measure to the optimal model Variant 4.0.1. (**C**) Differences in BIC to the optimal model Variant 4.0.1.

| Probe number | 1 | | 2 | | 3 | | 4 | | 5 | |
|---|---|---|---|---|---|---|---|---|---|---|
| Cleavage site (CS) | NSP4-NSP5 | | NSP5-NSP6 | | NSP6-NSP7 | | NSP7-NSP8 | | NSP8-NSP9 | |
| Main protease expression | + | – | + | – | + | – | + | – | + | – |

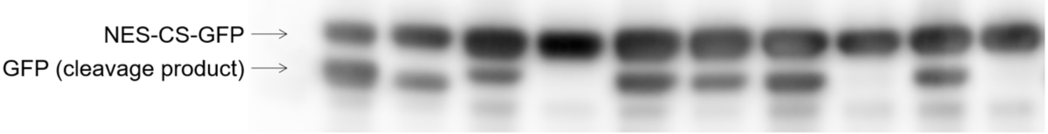


| Probe number | 6 | | 7 | | 8 | | 9 | | 10 | | 11 | |
|---|---|---|---|---|---|---|---|---|---|---|---|---|
| Cleavage site (CS) | NSP9-NSP10 | | NSP10-NSP11, NSP11-NSP12 | | NSP12-NSP13 | | NSP13-NSP14 | | NSP14-NSP15 | | NSP15-NSP16 | |
| Main protease expression | + | – | + | – | + | – | + | – | + | – | + | – |

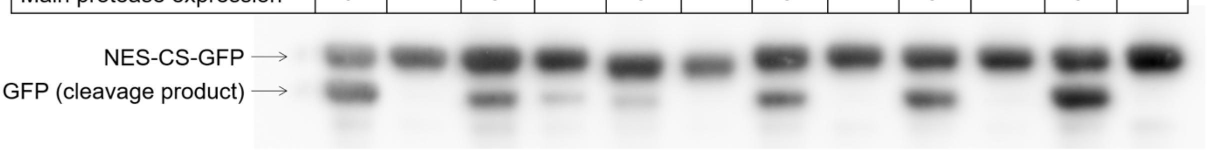


**Figure S8. Screening of 3CL$^{pro}$ cleavage probes by immunoblotting.** SARS-CoV-2 main protease (3CL$^{pro}$) cleavage sites in the precursor protein for non-structural proteins NSP4 to NSP16 were incorporated between a nuclear export sequence (NES) and GFP. Probe cleavage upon co-expression of 3CL$^{pro}$ and the respective probes was monitored using an anti-GFP antibody. For all probes, an increase in the band corresponding to the GFP part of the probe was detected upon 3CL$^{pro}$ co-expression, which is indicative of probe cleavage. Of note, a GFP-band was detected for probes number 1, 3 and 7 also in the absence of 3CL$^{pro}$. In these cases, cleavage motifs contained a methionine residue (Table S3) likely serving as alternative start codon resulting in expression of the GFP part independent of 3CL$^{pro}$ presence.

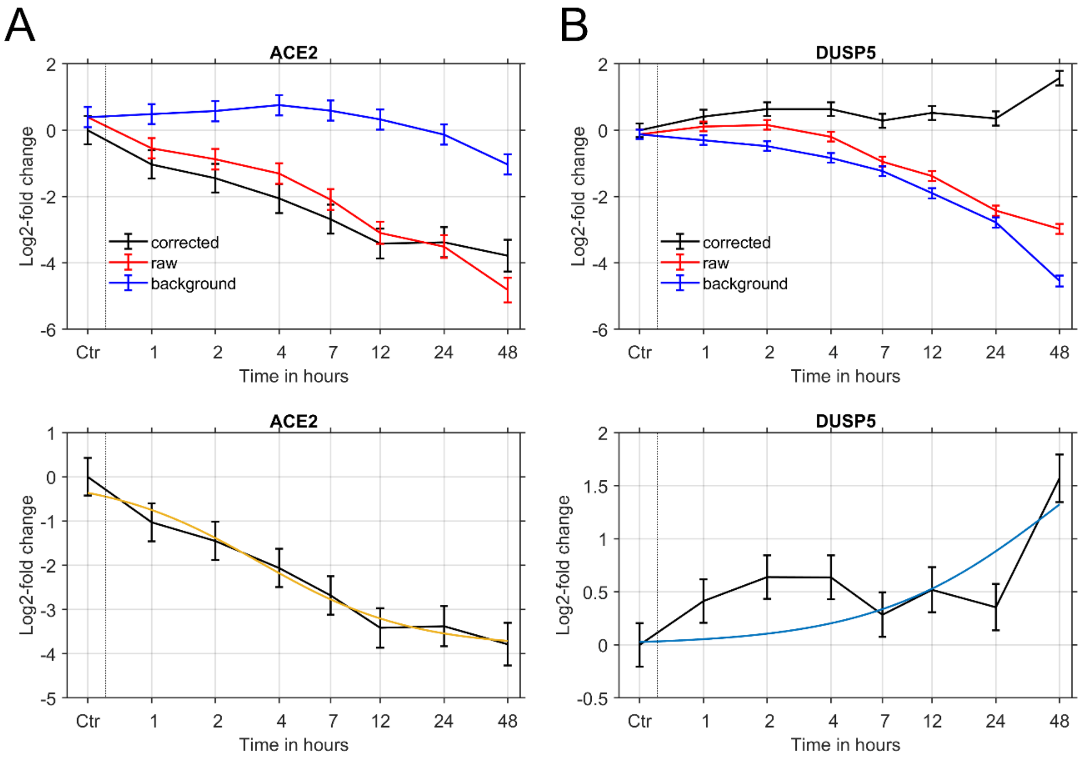


**Figure S9. Pre-processing of RNA-seq data.** (**A**, top) Raw values for from infected samples ('raw'), uninfected samples ('background') and differences ('corrected'). Error bars for measurements of infected samples indicate standard errors of triplicates, error bars for background values indicate values from the error model fitted to standard errors of mock samples, and error bars of corrected samples were inferred from error propagation. In case of ACE2, expression changes in mock samples were small. Therefore, values for corrected samples were similar to raw values from infected samples. (**A**, bottom) Corrected log2 fold changes for ACE2 and fitted profile function describing a continuous expression decrease (cf. Fig. 3). (**B**, top) Raw, background and corrected log2 fold changes for DUSP5 as in (A). In this gene, expression decreased over time in infected samples stronger than in mock samples. (**B**, bottom) Corrected log2 fold changes for DUSP5 and fitted profile function describing a continuous increase of expression (blue line).

#### Supplementary tables

**Table S1. Equations of the SARS-CoV-2 replication model variants.**

| Equation | Part of model version | Description |
|---|---|---|
| $\frac{d[V]}{dt} = k_{syn,V}[V]\frac{K_A}{K_A+[A]} - k_{deg,V}[V]$ | Variants 1, 4, 4.0.1–5, 4.3–5, 5, 6, 7 | Synthesis of virus transcripts $V$ inhibited by anti-viral proteins $A$ |
| $\frac{d[V]}{dt} = k_{syn,V}[V]\frac{K_A}{K_A+[A]}\frac{[P]}{K_P+[P]} - k_{deg,V}[V]$ | Variants 2, 4.1 | Synthesis of $V$ inhibited by $A$, accelerated by virus proteins $P$ |
| $\frac{d[V]}{dt} = k_{syn,V}[V]\frac{K_A}{K_A+[A]} - k_{deg,V}[V] + k_{influx,P}[P]$ | Variants 3, 4.2 | Synthesis of $V$, inh. by $A$, virus influx depending on $P$ |
| $\frac{d[m_A]}{dt} = k_{syn,m}\frac{[V]}{K_m+[V]} - k_{deg,m}[m_A]$ | Variants 1, 2, 3, 6 | Virus transcripts $V$ induce mRNAs of anti-viral genes $m_A$ |
| $\frac{d[m_A]}{dt} = k_{syn,m}\frac{[V]}{K_m+[V]}\frac{K_{inh,P}}{K_{inh,P}+[P]} - k_{deg,m}[m_A]$ | Variants 4, 4.0.1–5 4.1, 4.2, 4.4 | $V$ induce $m_A$ inhibited by $P$ |
| $\frac{d[m_A]}{dt} = k_{syn,m}\frac{[V]}{K_m+[V]}\frac{K_{inh,P}}{K_{inh,P}+[P]} - (k_{deg,m}+k_{cl,m}[P])[m_A]$ | Variant 4.3 | $P$ inhibit synthesis and induce degradation of $m_A$ |
| $\frac{d[m_A]}{dt} = k_{syn,m}\frac{[V]^h}{{K_m}^h+[V]^h}\frac{K_{inh,P,m}}{K_{inh,P,m}+[P]} - k_{deg,m}[m_A]$ | Variant 4.5 | Synthesis of $m_A$ depends on threshold of $V$ and is inhibited by $P$ |
| $\frac{d[m_A]}{dt} = k_{syn,m}\frac{[V]}{K_m+[V]} - (k_{deg,m}+k_{cl,m}[P])[m_A]$ | Variant 5 | $P$ induces degradation of $m_A$ |
| $\frac{d[m_A]}{dt} = k_{syn,m}\frac{[V]^h}{{K_m}^h+[V]^h} - k_{deg,m}[m_A]$ | Variant 7 | Synthesis of $m_A$ depends on threshold of $V$ |
| $\frac{d[A]}{dt} = k_{syn,A}\frac{[m_A]}{K_T+[m_A]} - k_{deg,A}[A]$ | Variants 1–5,7, 4.1–3, 4.5, 4.0.3 | Translation of $m_A$ to anti-viral proteins $A$ |
| $\frac{d[A]}{dt} = k_{syn,A}\frac{[m_A]}{K_T+[m_A]}\frac{K_{inh,P,A}}{K_{inh,P,A}+[P]} - k_{deg,A}[A]$ | Variants 6, 4.4 | Translation of $m_A$ to $A$ inhibited by $P$ |
| $\frac{d[A]}{dt} = k_{turnover,A}\frac{[m_A]}{K_T+[m_A]} - k_{turnover,A}[A]$ | Variants 4.0.1, 4.0.5 | Description of $m_A$ turnover by single parameter |

| | | |
|---|---|---|
| $$\frac{d[A]}{dt} = k_{syn,A}[m_A] - k_{deg,A}[A]$$ | Variants 4.0.2, 4.0.5 | Synthesis of $A$ by mass action instead of MM-kinetics |
| $$\frac{d[A]}{dt} = k_{turnover,A}[m_A] - k_{turnover,A}[A]$$ | Variant 4.0.4 | Mass action kinetics, single turnover parameter |
| $$\frac{d[P]}{dt} = k_{syn,P}\frac{[V]}{K_T + [V]} - k_{deg,P}[P]$$ | Variants 1–6, 4.1–5, 4.0.1, 4.0.2, 4.0.4, 4.0.5, 4.0.6 | Translation of virus transcripts $V$ to virus proteins $P$ |
| $$\frac{d[P]}{dt} = k_{syn,P}[V] - k_{deg,P}[P]$$ | Variants 4.0.3, 4.0.5 | Synthesis of $P$ by mass action instead of MM-kinetics |

**Table S2. Parameters of the optimal SARS-CoV-2 replication model 'Variant 4.0.1'.**

| Parameter | Unit | Best fit value | Mean | Std | VarK | 1σ-C.I., estimated by inverse of Hessian | | Allowed parameter interval | |
|---|---|---|---|---|---|---|---|---|---|
| | | | | | | Lower bound | Upper bound | Lower bound | Upper bound |
| $k_{syn,V}$ | $h^{-1}$ | 15.6 | 16.1 | 2.04 | 0.126 | 15.6 | 15.6 | $10^{-5}$ | 100 |
| $k_{deg,V}$ | $h^{-1}$ | 7.68 | 4.08 | 2.74 | 0.671 | 7.32 | 8.07 | $10^{-5}$ | 10 |
| $k_{syn,m}$ | $h^{-1}$ | 0.927 | 0.770 | 0.178 | 0.231 | 0.00949 | 90.5 | $10^{-5}$ | 1 |
| $K_{inh,P,m}$ | unitless | 0.0229 | 0.0348 | 0.0136 | 0.392 | $4.58\cdot10^{-5}$ | 11.4 | $10^{-5}$ | 1 |
| $k_{deg,m}$ | $h^{-1}$ | 0.0789 | 0.0869 | 0.00650 | 0.0747 | 0.0248 | 0.250 | $10^{-5}$ | 1 |
| $k_{turnover,A}$ | $h^{-1}$ | 0.412 | 0.360 | 0.0438 | 0.122 | 0.323 | 0.526 | $10^{-5}$ | 1 |
| $k_{syn,P}$ | $h^{-1}$ | 0.0431 | 0.0416 | 0.00252 | 0.0605 | 0.0277 | 0.0671 | $10^{-5}$ | 1 |
| $k_{deg,P}$ | $h^{-1}$ | 0.0428 | 0.0396 | 0.00566 | 0.143 | 0.0148 | 0.124 | $10^{-5}$ | 1 |
| $K_A$ | unitless | 0.959 | 0.428 | 0.391 | 0.913 | 0.958 | 0.960 | $10^{-5}$ | 1 |
| $\widetilde{K}_m{}^*$ | unitless | $3.74\cdot10^{-5}$ | $5.56\cdot10^{-5}$ | $4.31\cdot10^{-5}$ | 0.776 | $3.08\cdot10^{-8}$ | 0.0454 | $10^{-5}$ | 1 |
| $K_T$ | unitless | $4.04\cdot10^{-5}$ | $7.78\cdot10^{-5}$ | $9.94\cdot10^{-5}$ | 1.278 | $2.36\cdot10^{-8}$ | 0.0693 | $10^{-5}$ | 1 |
| $[V_0]$ | unitless | $1.11\cdot10^{-6}$ | $1.33\cdot10^{-6}$ | $6.54\cdot10^{-7}$ | 0.491 | $4.59\cdot10^{-7}$ | $2.69\cdot10^{-6}$ | $10^{-6}$ | 1 |

VarK, coefficient of variation; C.I., confidence interval; * the parameter $K_m$ was defined by $K_m = \widetilde{K}_m + [V_0]$, according to the assumption that virus replication was not saturated at the time of transfection.

**Table S3. Cleavage motifs of probes for monitoring main protease (3CL$^{pro}$).**

| Probe | Cleavage site | Cleavage motif |
|---|---|---|
| 1 | NSP4-NSP5 | SITSAVLQ‖SGFRKMAF |
| 2 | NSP5-NSP6 | QCSGVTFQ‖SAVKRTIK |
| 3 | NSP6-NSP7 | CIKVATVQ‖SKMSDVKC |
| 4 | NSP7-NSP8 | LDNRATLQ‖AIASEFSS |
| 5 | NSP8-NSP9 | ANSAVKLQ‖NNELSPVA |
| 6 | NSP9-NSP10 | LAATVRLQ‖AGNATEVP |
| 7 | NSP10-NSP11, NSP11-NSP12 | QLREPMLQ‖SADAQSFL |
| 8 | NSP12-NSP13 | YTPHTVLQ‖AVGACVLC |
| 9 | NSP13-NSP14 | RRNVATLQ‖AENVTGLF |
| 10 | NSP14-NSP15 | WNTFTRLQ‖SLENVAFN |
| 11 | NSP15-NSP16 | ETFYPKLQ‖SSQAWQPG |

Cleavage sequences were placed between an NES sequence and GFP. Cleavage sites are indicated by '‖' symbols.
